# Pericentromeric repeat copy number tunes heterochromatin dosage to control chromosome segregation and gene expression in fission yeast

**DOI:** 10.64898/2026.05.08.723760

**Authors:** Sarah E. Gilmour, Brandon L. Fagen, Devika Salim, María Angélica Bravo Núñez, Jeffrey J. Lange, Christopher Wood, Andrew Price, Michael T. Eickbush, R. Blake Billmyre, Alexandria J. Cockrell, Scott McCroskey, Martin Searcy, Kobe Koren, Luis Felipe Ramírez-Sánchez, Jennifer L. Gerton, Sarah E. Zanders

## Abstract

Centromeres are essential for chromosome segregation, yet in many genomes they are composed entirely of rapidly evolving repetitive DNA, embedded in other repetitive DNA that forms pericentromeric heterochromatin. Due to the difficulties of manipulating these repeat-rich regions, how the relative size of pericentromeric repeat regions influences chromosome segregation remains an open question. Here, we take advantage of the tractable *Schizosaccharomyces pombe* system by combining population-level analysis, complete long-read assemblies, and engineered near-isogenic strains to test how pericentromeric repeat copy number affects chromosome biology in its native context. We find that pericentromeric *dh/dg* arrays on chromosome 3 vary almost tenfold in size among natural *S. pombe* isolates, ranging from 35 to 265 kb. We converted this natural diversity into an experimental system of nearly isogenic strains that primarily differ in pericentromere size (35 to >350 kb). We found that pericentromere size does not alter baseline growth under standard conditions. However, larger pericentromeres alter transcriptional output and sensitize cells to spindle stress. We show that this spindle-stress phenotype depends on heterochromatin: loss of the H3K9 methyltransferase Clr4 abolishes size-dependent differences, whereas artificial targeting of the Chromosomal Passenger Complex to heterochromatin partially rescues the defect. Thus, we find that larger pericentromeres act as sinks for limiting regulatory factors, weakening their effective concentration at centromeres and compromising faithful chromosome segregation under stress. These results establish that naturally occurring copy-number variation within repetitive pericentromeric DNA is not merely noise, but a functional source of variation in chromosome segregation and gene regulation. Our work provides an experimentally tractable framework for understanding how repeat expansion in centromere-proximal heterochromatin influences chromosome behavior across eukaryotes.

## Introduction

Centromeres are regions of chromosomes where kinetochores assemble and spindles attach during cell division. Centromeres are not defined by a specific DNA sequence but are usually defined epigenetically by the deposition of a centromeric histone H3 variant and heterochromatinization of adjacent pericentromeric sequences (1,2). In most eukaryotes, centromeres and pericentromeres consist of long arrays of repetitive sequences (3–5). Copy number variation in these repetitive sequences is a major source of interindividual variation in humans and in mice, with the size of an array varying up to 50-fold (6–9). The functional consequences of this copy number variation are unknown because these arrays are genetically intractable, even in model systems. In this study, we overcome this technological barrier using the fission yeast *Schizosaccharomyces pombe* to address whether and how naturally occurring variation in the size of repetitive centromere-proximal DNA affects cell fitness and function.

Studies of *S. pombe* centromeres have helped to establish many critical aspects of centromere biology, such as the role of chromatin modification in supporting centromere identity and function (10–13). *S. pombe* centromeres are composed of a central core domain (*cc*) flanked by the innermost repeats (*imrs*), which together serve as the site of kinetochore assembly (14). The centromere is flanked by heterochromatic pericentromeric arrays, composed of *dh* and *dg* repeats (15–20). The use of distinct sequences for kinetochore assembly and pericentromeres in *S. pombe* facilitates disambiguation of core and pericentromere functions.

The heterochromatin of the *dh/dg* repeat arrays (14,16) facilitates chromosome segregation by promoting sister chromatid cohesion (10,11,16,21–24),and by recruiting essential cell cycle regulatory proteins, such as the Chromosomal Passenger Complex (CPC) which regulates mitotic progression (25–29). Beyond the centromere, it is possible that the amount of pericentromeric heterochromatin could have broad effects on the global chromatin landscape by acting as a sink for various heterochromatin factors (30). In *Drosophila*, for example, distinct Y chromosomes with differing amounts of heterochromatin alter genome-wide distribution of heterochromatin and gene expression (31,32).

Here we surveyed extant copy number variation in 160 *S. pombe* strains and assembled the centromere regions of seven strains. We observed a striking degree of *dh/dg* copy number variation on chromosome 3, with total size ranging from 35 to ∼265 kb. We then leveraged this variation to generate the first set of near-isogenic strains that specifically vary tenfold in the size of the repetitive centromere region (from 35 kb to >350 kb) on one chromosome. We find that enlarged pericentromeres change transcriptional output and selectively impair growth under spindle stress. Second, we show that this size-dependent spindle-stress phenotype requires heterochromatin and can be partially rescued by artificially targeting the Chromosomal Passenger Complex to heterochromatin. Our study thus links repeat expansion directly to chromosome segregation phenotypes and supports a model in which expanded pericentromeric heterochromatin acts as a sink to dilute limiting regulatory factors.

## Results

### Natural variation in in *S. pombe* centromeres

To broadly survey pericentromeric copy number variation, we used a droplet digital PCR (ddPCR) approach to quantify the total copy number of *dh* and *dg* in a collection of 160 natural isolates (Table S4) (33). We found a wide distribution of copy numbers, as low as ∼7 and as high as ∼38 (Fig S1A-B), in line with previous findings (17,19). Consistent with rapid evolution of centromere sequences, we found no correlation between *dh*/*dg* copy numbers and phylogenies of these natural isolates (Fig S1C-D).

We then assembled the centromeres in seven *S. pombe* strains using Nanopore long-read sequencing to understand the variation at higher resolution. The strains we chose span the known diversity of *S. pombe* and include two lab strains (SZY44 and SPT999a) clonally derived from the same background (Fig 1A) (33–37). We confirmed the quality of our assembly of the lab strains (SZY44 and SPT999a) by comparing them to the published reference genome and found that the assemblies were highly identical. In six of the seven strains, we obtained a complete assembly. We failed to fully assemble the long *dh/dg* repeat array on the left side of centromere 3 in JB907. However, we did fully assemble the same pericentromeric array introgressed into another strain (“Large” strain, Fig 2A) so we used that assembly for centromere 3 only of JB907.

**Figure 1.**
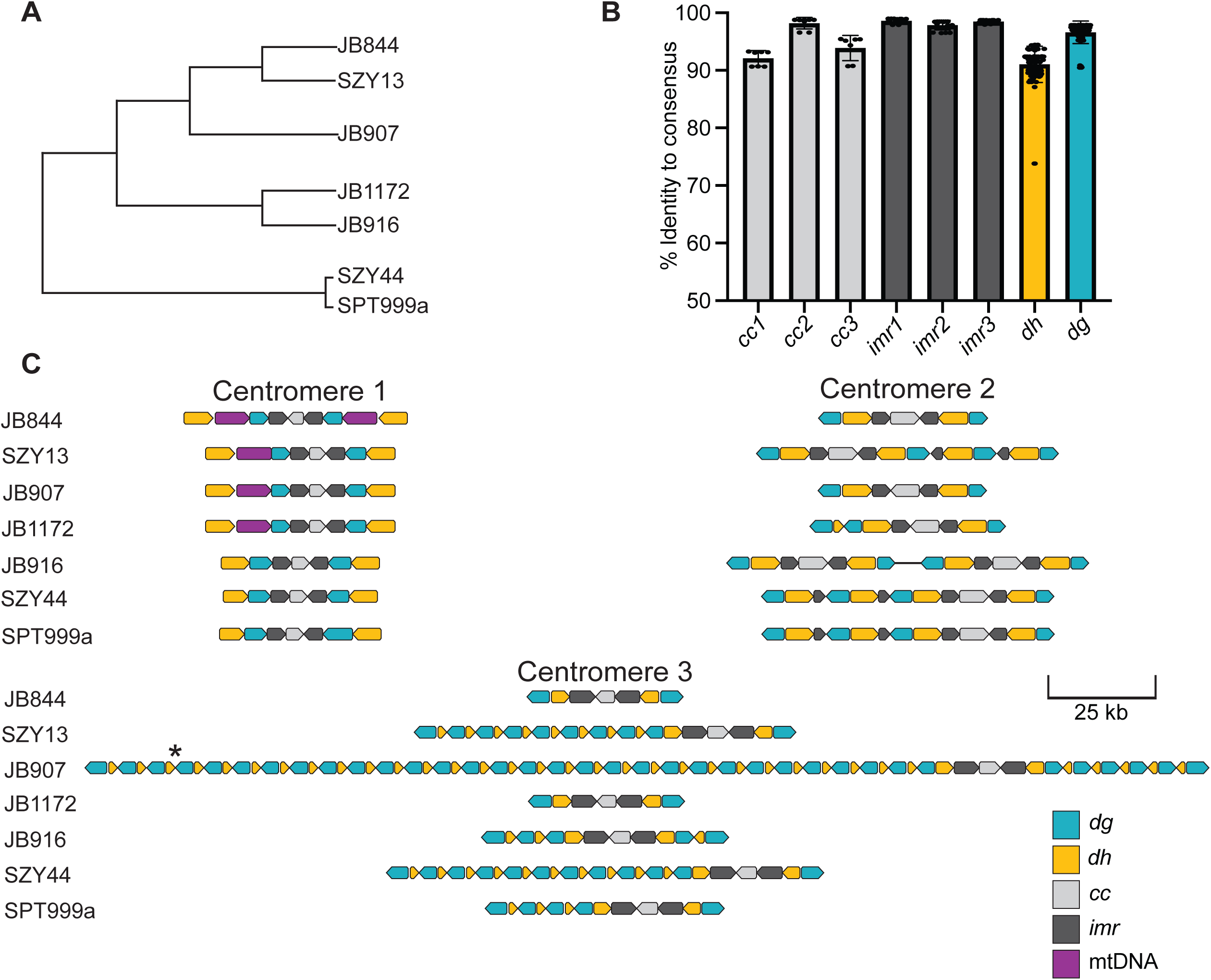
Extensive natural variation exists in *S. pombe* centromeres. A) Phylogeny of natural isolates assembled in this study. The tree topology shown is derived from (65). SZY13 is derived from the *S. kambucha* isolate, which is highly similar to JB1180. SZY44 and SPT999a are derived from Leupold’s 972 reference strain, which is highly similar to JB22. B) Each centromere sequence element from the assemblies is compared to a consensus sequence derived from all the sequences. The percent sequence identity of each element to the consensus was plotted, with each dot representing one sequence. Raw data are available in Table S5. C) Graphical representation of sequence elements based on long read sequencing assemblies of complete centromeric regions. The elements on all 3 chromosomes are drawn to scale. *The assembly of centromere 3 from JB907 contained a gap, so the corresponding SZY5827 assembly is shown. Different assembly techniques yielded either 30 or 35 copies of *dh/dg* in the left side of the array, and sequencing coverage estimates (Table S6) further support that size range.

**Figure 2.**
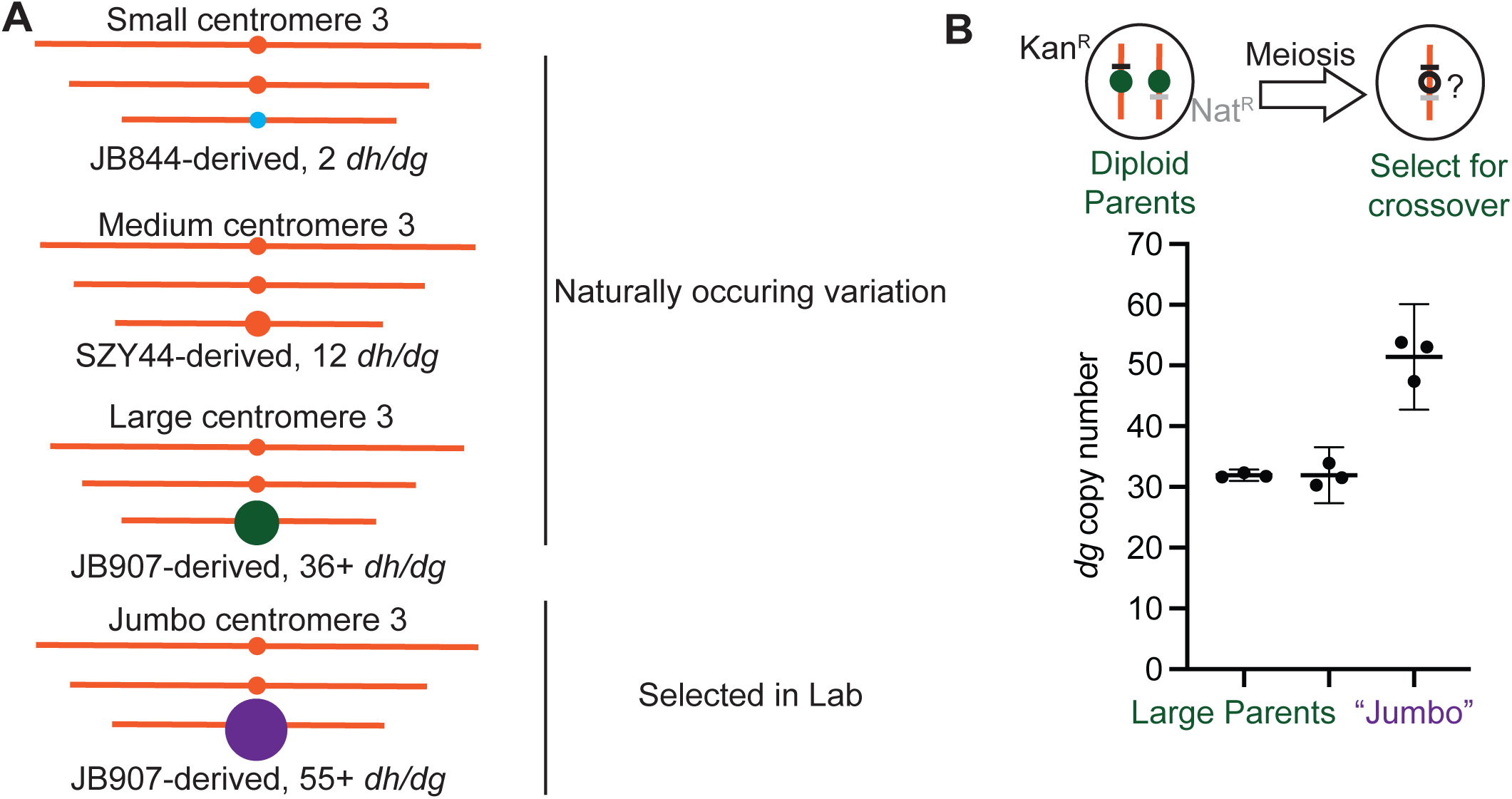
Generating isogenic centromere 3 variant strains. A) Schematic depiction of the Small, Medium, Large and Jumbo centromere 3 alleles used in this study. All strains share the lab isolate (SZY44) strain background (see Fig S4 for details). B) Schematic depiction of the selection method for finding centromere-proximal crossovers (top). Diploids with different drug markers on either side of the centromere were put through meiosis, and haploid progeny with resistance to both markers (Nat^R^ Kan^R^) were selected and measured via ddPCR. The ddPCR results of the haploid parents and the most substantially altered offspring, termed “Jumbo” are shown on the right. ddPCR was performed in technical triplicate. All 3 technical replicates are plotted, along with the median values (lines) and replicates are available in Table S13.

Consistent with rapid evolution of centromere sequences, we found the sequence divergence in centromeres and pericentromeres between isolates (3-8%) is higher than what is observed between the genome at large (<1%) (Fig 1B, Fig S2-S3, Table S5) (37). On chromosome 1, we found no variation in *dh/dg* copy number, consistent with previous work (17). However, several natural isolates have a large insertion (8.1 kb) of mitochondrial DNA (Fig 1C) (34). On chromosome 2, we observed 2-4 copies of the *dh* and *dg* repeats, similar to previous observations (Fig 1C) (17). However, one isolate (JB916), contains a novel centromere structure derived from a duplication of the centromere and pericentromere on chromosome 2 (Fig 1C).

On chromosome 3, the smallest centromere has 2 copies of *dh/dg* and is about 37 kb in length while the largest has at least 36 *dh/dg* repeats and is at least 265 kb in length, expanding previous estimates for the largest *S. pombe* centromere region by >100 kb (17,19). The exact size of the larger pericentric array varies slightly based on assembly parameters and may be as large as 300 kb, with sequencing coverage supporting either of those sizes (Table S6). Given the small size of chromosome 3 (∼3.5 Mb), this centromere size variation drives significant variation in overall chromosome size.

### Building centromere-variant isogenic strains

Identifying potential functional consequences of repeat variation requires separating copy number-dependent phenotypes from other effects of genetic background. We achieved this by swapping a “Small” (37 kb, from JB844), “Medium” (100 kb, from SZY44), and “Large” (∼265 kb, from JB907) centromere 3 allele into the genetic background of the lab strain SZY44 (Fig 2A). The core centromeres in these strains are highly similar, sharing 95.4% identity across the *cc*, *imrs*, and first *dh/dg* repeat on either side of the *imrs* (Fig S3C).

To build the “swap” strains, we used a series of crosses and targeted recombination (Cre/lox) events (35) (Figs 2A, S4). We assayed isogeneity of the strains using a combination of long and short read sequencing. We found 149 variants specific to the Large pericentromere swap strain (Table S10), only 6 of which fell within coding regions, 5 of which are nonsynonymous (Fig S5, Table S7). The Medium pericentromere swap strain similarly contained 162 unique variants (Table S9), with 4 falling inside coding regions, all nonsynonymous (Fig S5, Table S7). The Small pericentromere swap strain contained more variants, with 1116 unique variants (Table S8), including 62 coding region mutants. However, 59/62 of those variants are found within eight genes which are located between centromere 3 and the location that the chromosome arm “swap” occurred. There was only one other gene with coding sequence variation in the Small pericentromere strain (Fig S5, Table S7). By contrast, distinct natural isolates of *S. pombe* can have roughly 1 SNP per 200 base pairs, including in coding regions (33,36). Our sequencing also revealed that three *dh/dg* repeats flanking centromere 2 had collapsed into one repeat in the common ancestor of the swap strains (Fig 1C, S2).

All the strains used in subsequent experiments were derived from the sequenced strain set by crossing and/or transformation and are thus expected to maintain similarly high levels of isogeneity. Moreover, the pericentromere sizes were stable through mitosis (Fig S6A, Table S14) and during strain construction crosses.

### Rare centromere-proximal crossovers can produce new copy number alleles

In addition to studying the largest and smallest naturally occurring centromere 3 alleles, we wanted to expand our study into additional pericentromere sizes. We were also curious how pericentromeres with different copy numbers arise in the population. We hypothesized that rare, uneven meiotic crossovers in the *dh/dg* array could change copy numbers. Despite the pericentromere being a cold spot for meiotic recombination (37), meiotic crossovers within the pericentric region of *S. pombe* do occur (12).

To assess meiotic recombination at pericentromere 3, we employed heterozygous markers (*Nat^R^* and *Kan^R^*) directly adjacent to the centromeric array on either side (Fig 2B). We recovered recombinants at a very low rate of about 0.3% independent of centromere size (Fig S7A, Table S12). Out of 270 recombinant progeny screened via ddPCR, we discovered one with a dramatically higher *dh/dg* copy number. We were unable to assemble this novel allele, which we termed “Jumbo”, but ddPCR and sequence read depth allow us to estimate there are ∼50-55 copies leading to a centromere region of ∼350-380 kb (Fig 2B, Table S6, Table S13). The discovery of Jumbo confirms that copy number variation can arise during very rare pericentromeric crossover events and offered an opportunity to assess the functional consequences of an extreme pericentromere size.

Like the other swap strains, we observed few unique variants (283 total, 35 non-synonymous) in the Jumbo genome (Table S11). However, the reads used to generate this assembly were shorter than the reads used in the other assemblies (N50 of 11 kb for Jumbo vs 28-30 kb for Small, Medium, and Large) and the telomeres were not efficiently assembled. While fully assembled *S. pombe* telomeres are expected to include 4 copies of the *tlh* genes (38), inadequate telomere assemblies, such as in our Jumbo assembly, often consolidate the *tlh* genes into fewer than 4 copies. Accordingly, 60/283 (21%) of all variants and 12/35 (34%) of nonsynonymous variants were found within 20 kb of the chromosome ends (Table S7). (41)

### Pericentromere size affects global gene expression

To test if pericentromere size globally affects transcription, we sequenced mRNA isolated from swap strain cells in logarithmic growth under standard conditions. We compared the swap strains to the Medium pericentromere strain, as that allele is endogenous to the SZY44 strain background shared by all the swap strains. Note that this experiment used strains that contained a tagged histone (*hht1-RFP*), as this allele likely served as an enhancer of at least one phenotype (see below).

The Small and Large strains have only 4 and 8 genes, respectively, that are differentially expressed relative to the Medium strain (Table S15). Notably, nearly all differentially expressed genes are long non-coding RNAs or pseudogenes which fall within constitutive heterochromatin in the pericentromeres or subtelomeres (Fig 3A-B) (39). Subtelomeric transcripts are downregulated in the Small strain and upregulated in the Large strain. These patterns are consistent with less heterochromatin at pericentromere leading to increased silencing at heterochromatin elsewhere, while more heterochromatin at the pericentromere leads to decreased silencing at heterochromatin elsewhere.

**Figure 3.**
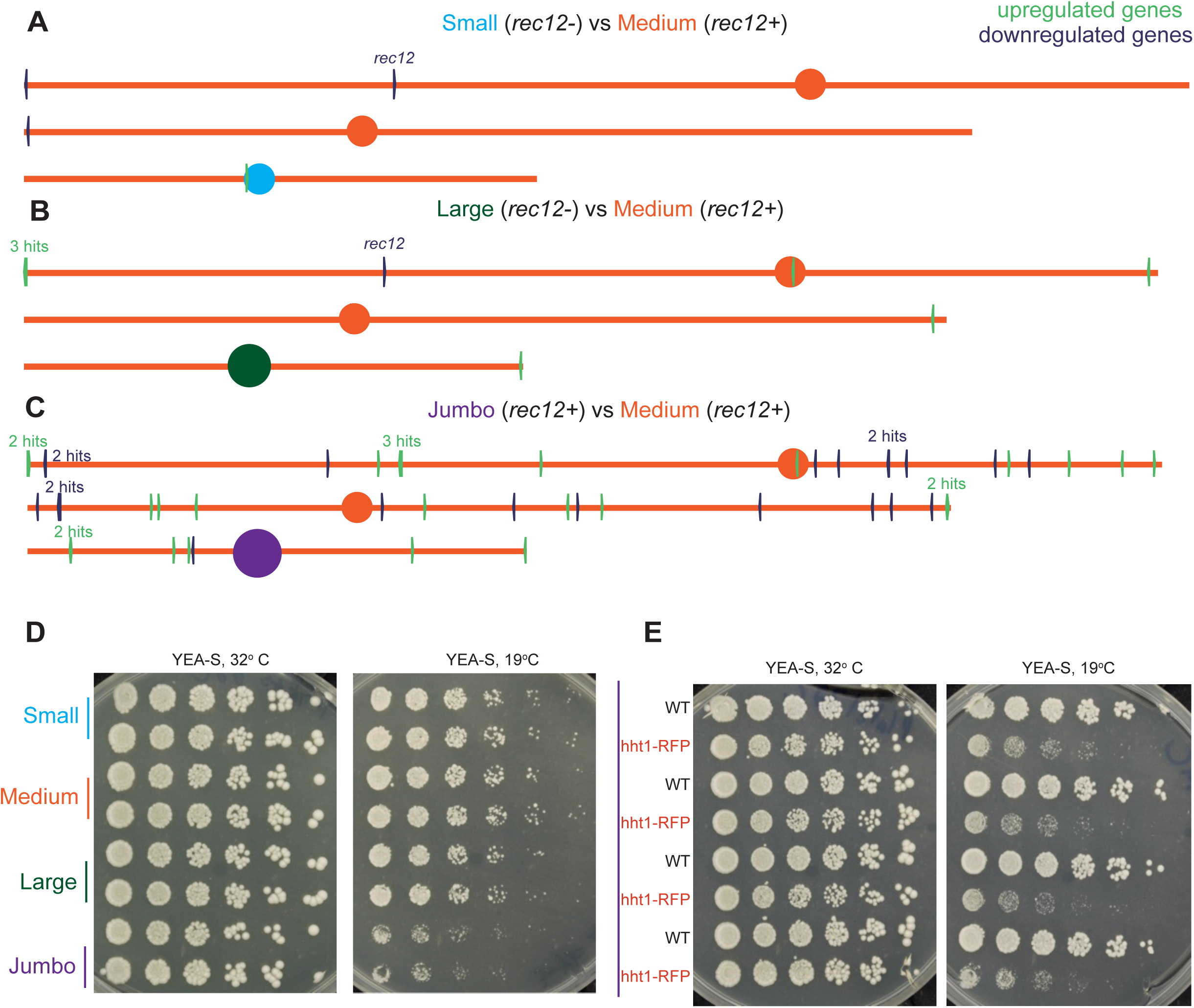
Differentially expressed genes correlate with constitutive heterochromatin. A-C) The genomic locations of all differentially expressed genes (listed in Table S15) are shown, with upregulated genes in green and downregulated genes in navy. D) Spot assays are shown of Small, Medium, Large, and Jumbo centromere 3 strains (2 biological replicates shown, all *hht1-RFP*) onto the indicated media. The plates were incubated for three days at 32°C or six days at 19°C prior to imaging. E) Spot assays of *hht1-RFP* and WT Jumbo strains, meiotic products of a WT/*hht1*-*RFP* heterozygous diploid, are shown on indicated media. The synthetic effect of the Jumbo allele and the tagged histone yield cold sensitivity.

The Jumbo strain exhibits more dramatic transcriptional changes, with 45 differentially expressed transcripts, including long non-coding RNAs and pseudogenes (Fig 3C, Table S15). Amongst the 14 coding genes with increased transcripts in the Jumbo strain is *mug170*, a gene whose transcription is normally upregulated in meiosis (39,40). The telomere helicases *tlh1* and *tlh2*, which are encoded in subtelomeres and are overexpressed during telomere crisis or in mutants that disrupt subtelomeric heterochromatin, also have higher transcript levels in the Jumbo strain (41,42). A final noteworthy gene with increased transcripts in the Jumbo strain is *hsp16*, a heat shock protein folding chaperone whose transcripts increase during a variety of cellular stresses (39,43). The upregulation of these transcripts could suggest that this strain is undergoing stress related to loss of constitutive heterochromatin.

There are also 12 coding genes which have fewer transcripts in the Jumbo strain. These include six genes promoting iron import, which constitute the Gene Ontology classes (related to iron metabolism) significantly altered in our comparisons. These six genes are all bound and repressed by the histone deacetylase Clr6 (HDAC1) in cells grown in normal lab conditions (44), suggesting possible shifts in the usage of different heterochromatin pathways in this strain. Interestingly, iron is important for establishing facultative heterochromatin that facilitates growth at a low temperature and *hht1-RFP* Jumbo strains are cold-sensitive (Fig 3D). This is a synthetic phenotype requiring both the Jumbo centromere and *hht1-RFP* as Jumbo strains without *hht1-RFP* are not cold sensitive (Fig 3E).

The transcriptional changes seen in the Jumbo strain indicate that extreme pericentromeric array size can have a complex effect on the gene regulatory landscape and associated phenotypes. Taken together, our transcriptional analyses suggest that the size of a pericentromeric array can affect the distribution of heterochromatin across the genome.

### Larger pericentromere strains are sensitive to inhibition of microtubule polymerization

The centromere swap strains grew identically under standard growth conditions and in response to several stress conditions (Fig S6B). We similarly found no differences in chromosome segregation fidelity in unstressed meiosis (Fig S8, Table S16). The Large and Jumbo strains, however, are increasingly sensitive to the microtubule polymerization inhibitors Thiabendazole (TBZ) and benomyl (Fig 4A). These phenotypes are independent of the *hht1-RFP* marker (Fig 4A, Fig S5). As expected, TBZ-treated cells struggle to divide efficiently, often fail to form a proper septum, exhibit lagging chromosomes in anaphase, and segregate chromatin unevenly (Fig 4B). In concordance with the growth defects, the Large and Jumbo strains exhibit a higher frequency of unequal chromatin division with TBZ treatment, relative to the Small and Medium strains (Fig 4C, Table S17). The heterogeneity in the unequal chromatin masses suggests the missegregation impacts all chromosomes, not just chromosome 3. Our results indicate that TBZ induces cell death via a global increase in chromosome missegregation, and these events increase with pericentromere size.

**Figure 4.**
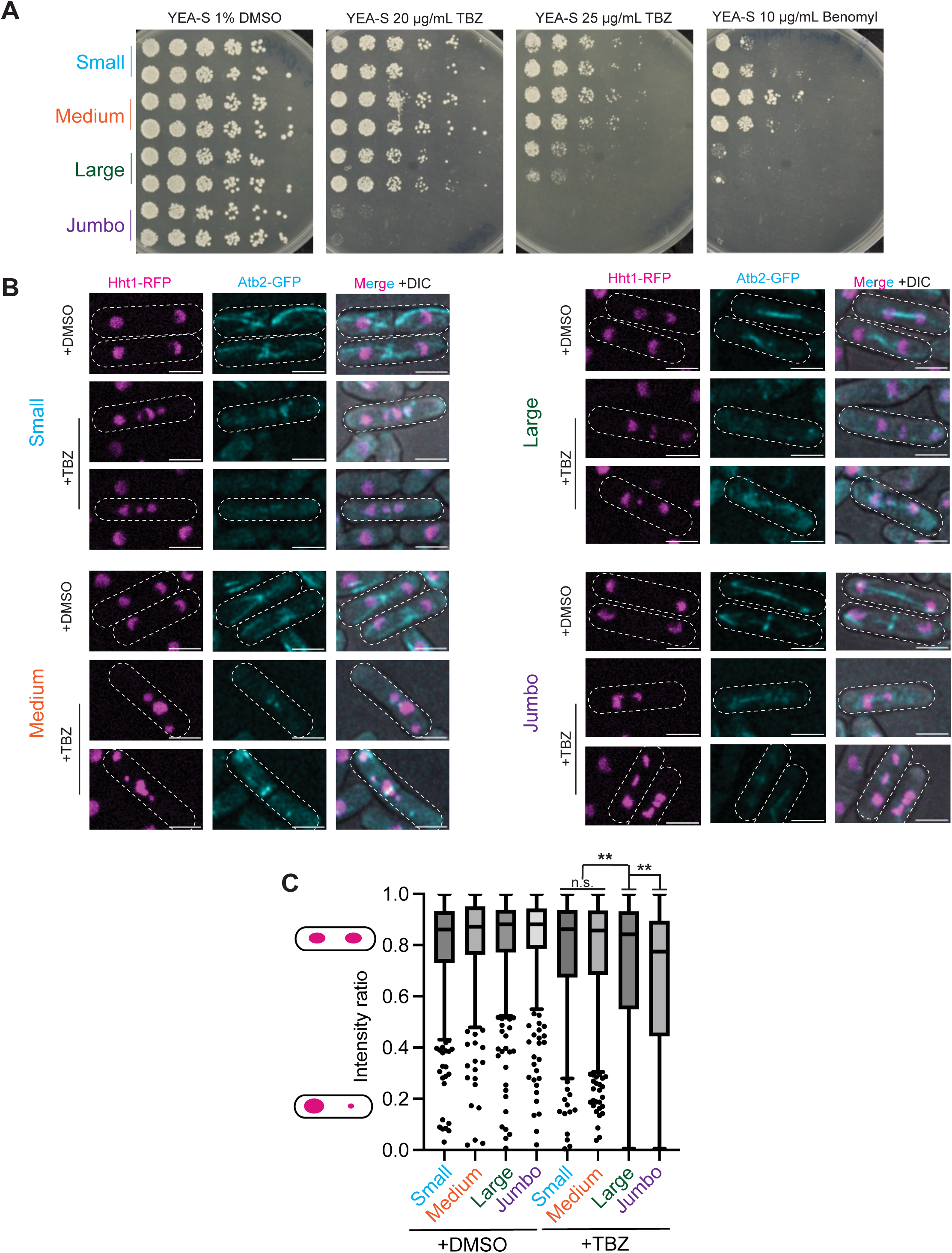
Larger centromere 3 strains are sensitive to spindle stress. A) Spot assays of Small, Medium, Large, and Jumbo cen3 strains (2 biological replicates) were grown under conditions of spindle stress (TBZ and Benomyl). All plates were incubated for 3 days at 32°C prior to imaging. B) Still images from timelapse microscopy of cells treated with DMSO or TBZ are shown. Cells expressing Hht1-RFP and Atb2-GFP were imaged every 20 minutes at 32°C on agar pads of YEA-S +1% DMSO or YEA-S + 25 μg/mL TBZ. Scale bar = 5μm C) Intensity ratio of Hht1-RFP foci in freshly divided cells is depicted. Each dot represents one pair of Hht1-RFP foci, with the intensity of the dimmer focus divided by the intensity of the brighter focus.Tukey boxplot is shown, with the central line of each box representing the median, the box representing the upper and lower quartile, and individually plotted points representing the outliers. n>290 pairs for each condition. Full counts are available in Table S17. ** indicates p<0.01 by Welch’s unpaired t-test.

### Size-specific phenotypes depend on heterochromatin, but not silencing

We predicted that the TBZ sensitivity differences between the strains stemmed from differences in pericentromeric heterochromatin on *dh* and *dg* repeats (45–47). To test this hypothesis, we deleted the histone methyltransferase gene *clr4* (*Suv39*), which is necessary to form all heterochromatin in *S. pombe* (48,49). *clr4* mutants have high sensitivity to TBZ and other spindle stressors (10,24). Importantly, the *clr4* mutant strains are equally sensitive to TBZ, indicating the pericentromere size-dependent phenotype requires heterochromatin (Fig 5A).

**Figure 5.**
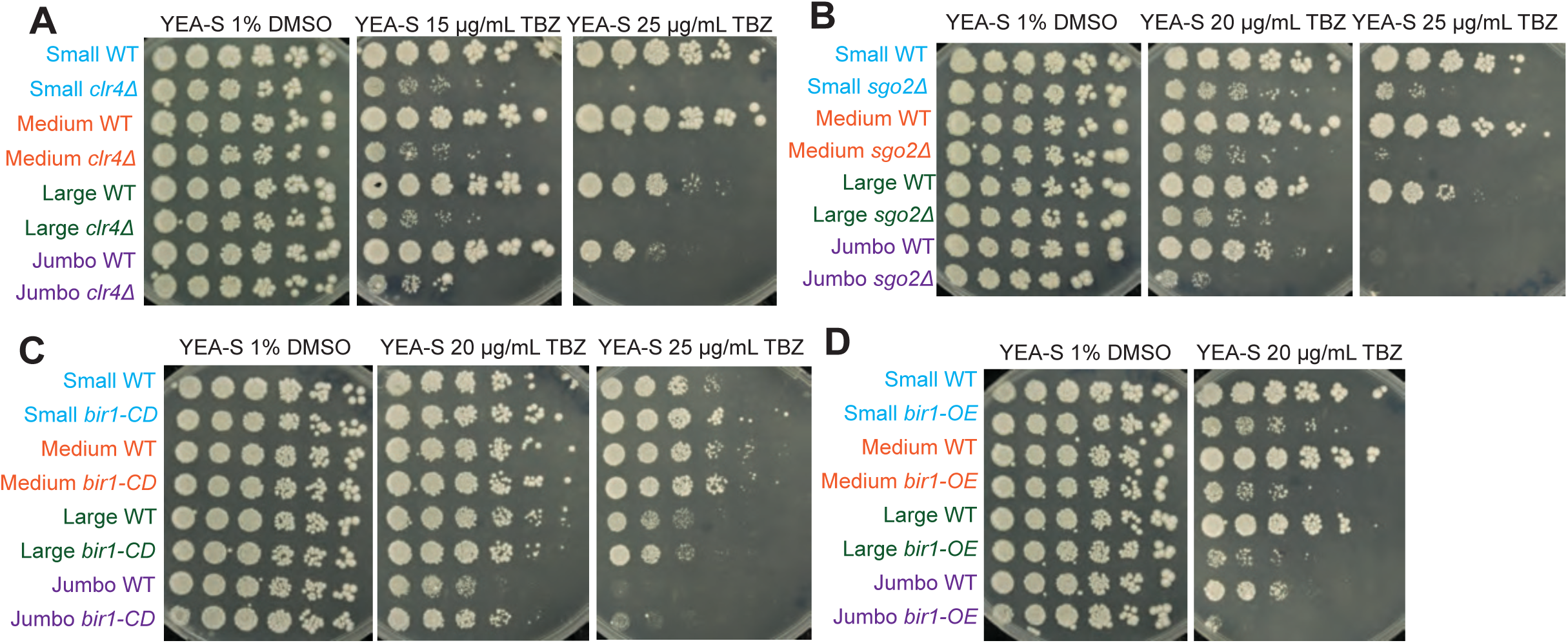
Heterochromatin and CPC factors modulate sensitivity to spindle stress. A-D) Spot assays of strains with the indicated genotypes and size alleles were grown on YEA-S at 32°C for 3 days with variable TBZ concentrations.

Pericentromeric heterochromatin both limits transcription and helps recruitment of many factors to the centromere region (21–23,25,50). To distinguish between these roles, we used an allele of the heterochromatin protein *swi6* (*HP1*)*, swi6-sm1,* which is deficient in biomolecular condensation and silencing but sufficient to recruit cohesin and shugoshin to the pericentromere (23). We found the *swi6-sm1* mutation had no effect on the TBZ sensitivity of our strains (Fig S9A). This suggests that the recruitment of other factors to heterochromatin, not differential condensation or transcriptional silencing, underlies pericentromere size-specific sensitivity to spindle stress.

### Copy-number dependent TBZ sensitivity is modulated by CPC factors

We hypothesized that the heterochromatin dependence of TBZ sensitivity may be due to altered recruitment of the Chromosomal Passenger Complex (CPC) to heterochromatin (29,51). The CPC is recruited to the pericentromere via two pathways, one that is heterochromatin-dependent and one that is not (52–54). Both pathways recruit the CPC via interactions with CPC component Bir1 (23,53).

In the heterochromatin-independent CPC recruitment pathway, the spindle checkpoint kinase Bub1 phosphorylates histone H2A-S121, which binds the shugoshin protein Sgo2, which then recruits the CPC to the pericentromere via interaction with Bir1 (28,55). We disrupted this pathway by deletion of *sgo2* (29), increasing TBZ sensitivity in all strains (Fig 5B). Importantly, TBZ sensitivity increases with pericentromere size (Fig 5B). As *sgo2Δ* strains are solely reliant on the heterochromatin-dependent CPC recruitment pathway, these results are consistent with our hypothesis that larger pericentromeres challenge the efficacy of heterochromatin-dependent CPC recruitment.

To further test our hypothesis, we attempted to rescue the sensitivity of the larger strains to TBZ by artificially tethering the CPC to pericentric heterochromatin. We achieved this by introducing an additional allele of *bir1* which is fused with a chromodomain (CD), which binds methylated H3K9 in heterochromatin (55). We observed that *bir1-CD* reduces the TBZ sensitivity for all pericentromere size strains, although the largest pericentromere strains remained the most sensitive (Fig 5C). Importantly, an additional allele of *bir1* without a chromodomain has the opposite effect in that it enhances sensitivity in all backgrounds (Figs 5D, S9B). We speculate this could be due to a change in stoichiometry of CPC components which could result in fewer complete CPCs being targeted to the pericentromere. Together, our results are consistent with inefficient enrichment of CPC at large pericentromeres contributing to size-dependent TBZ sensitivity.

We conclude that spindle stress sensitivity in strains with large pericentromeres is likely driven by an inability to regulate mitotic progression. While pericentromeric heterochromatin is essential to support correct chromosome segregation, an abundance of heterochromatin at just one centromere can strain the mechanisms that enrich key regulatory factors within the pericentric region, resulting in chromosome missegregation and cell death under spindle stress.

## Discussion

In this study, we describe the natural variation found within a broad sample of *S. pombe* strains. We identified extensive natural variation in pericentromere size, a natural chromosome 2 with two potential kinetochore assembly sites, and an allele of centromere 1 that surprisingly contains ∼40% of the mitochondrial genome. We then exploited the dramatic natural variation in chromosome 3 pericentromeric repeat copy number to engineer the first near-isogenic organisms that differ primarily in pericentromere size. This unique genetic advance allowed us to assess the biological effects of variation in pericentromere size at an organismal level for the first time.

We found that pericentromere size variation within the range observed in nature (Small to Large) affects global gene expression such that strains with a Small pericentromere have enhanced suppression of a set of genes, while strains with a Large pericentromere struggle to suppress expression of genes within heterochromatic regions. Extreme pericentromere size, represented by Jumbo, leads to more dramatic changes in the transcriptional landscape, including suppression a set of genes repressed by the histone deacetylase Clr6 (HDAC1) and overexpression of telomeric helicases (42,44). Finally, we found that chromosome segregation is disrupted in strains with Large and Jumbo pericentromeres under conditions of spindle stress (Fig 4).

Our data support a model in which pericentromeres act as a sink for heterochromatin-associated factors, wherein the pericentromere size of one centromere has global consequences for transcription (Fig 6). This model is supported by the altered transcription of genes encoded within constitutive heterochromatin (Fig 3) in the swap strains. We also propose expanded pericentromere size on one chromosome can compromise centromere function at least in part due to insufficient recruitment of the CPC to pericentromeres. This model is supported by the pericentromere size-dependent sensitivity to spindle stress and the phenotype’s dependence on heterochromatin. The suppression of the sensitivity to spindle stress via the addition of a chromodomain-linked copy of CPC factor *bir1* is also consistent with our model (Fig 5).

**Figure 6.**
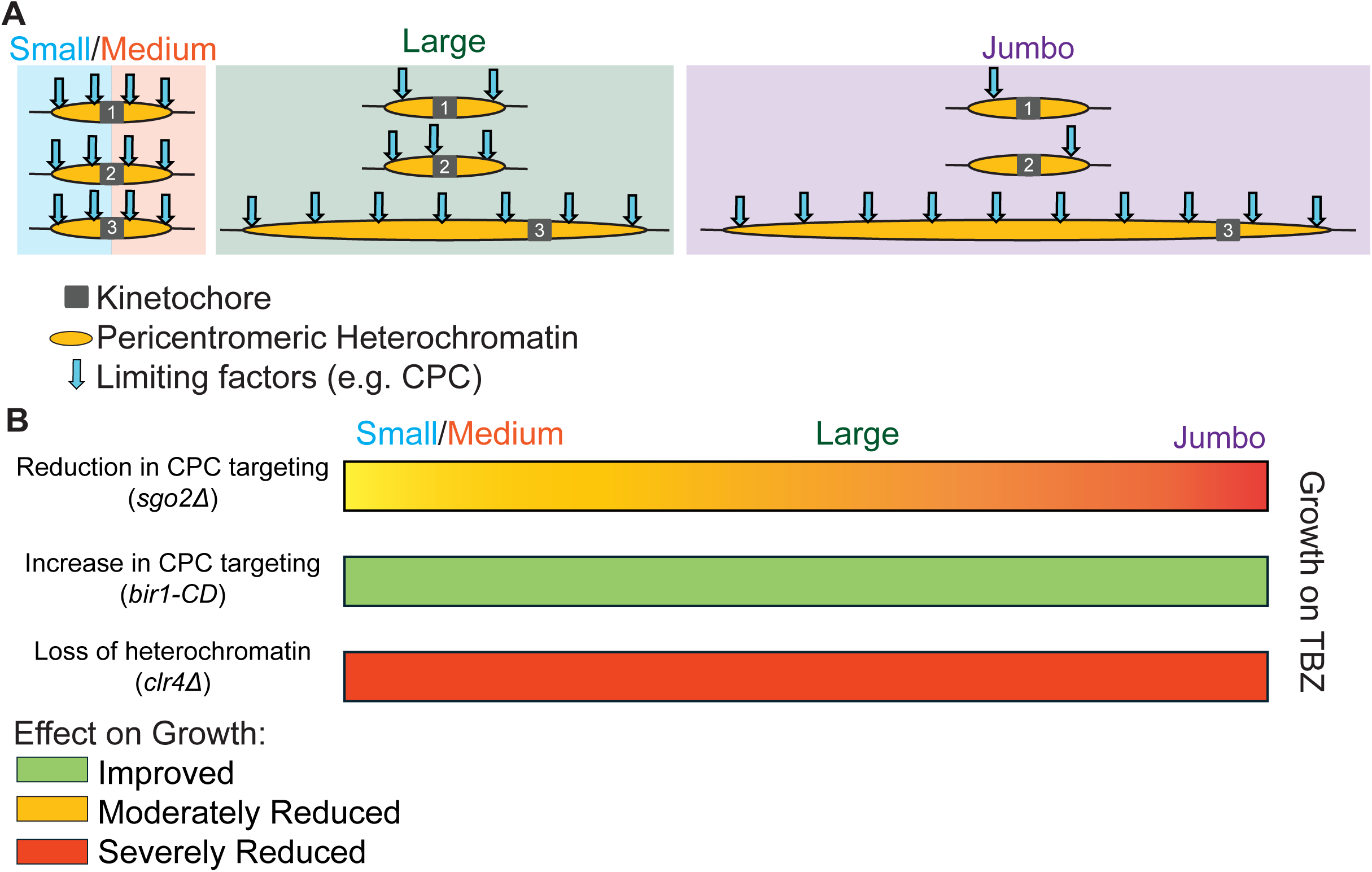
Working model for pericentromere size-dependent CPC function. A) The larger pericentromeres challenge CPC recruitment genome-wide due to the enlarged heterochromatin arrays acting as sinks for CPC proteins and other limiting factors. B) Summary of the effects of various types of mutations on TBZ-treated growth in strains with different pericentromere sizes. Loss of CPC targeting decreased growth on TBZ in all strains but more dramatically in strains with larger pericentromeres. Increasing CPC targeting improved growth on TBZ for all strains. Loss of heterochromatin decreased growth on TBZ for all strains in an equal fashion.

While our study focused on one pericentromere of *S. pombe*, copy number variation is prevalent across eukaryotes in many different types of heterochromatic repeats. Improved genome assemblies enabled by long read sequencing are continually expanding our understanding of this variation, which is already known to be a major factor distinguishing individual genomes within a species. Our work suggests this variation is likely to have global biological consequences.

## Materials and Methods

### Droplet digital PCR (ddPCR) for *dh* and *dg* copy number measurement

ddPCR is a quantitative method wherein PCR reactions are split into thousands of droplets and amplified with fluorescent probes to enable quantification of DNA molecules in the original sample relative to a single copy control gene. We isolated genomic DNA for ddPCR using the YeaStar Genomic DNA Kit (Zymo Research), measured concentrations on a Qubit Fluorometer and diluted the DNA to a concentration of 0.01 ng/uL in water. We used the manufacturer’s protocol to carry out duplexed ddPCR on the BioRad QX200 system. Briefly, we assemble master mixes including primers and probes for the target sequence (*dh* or *dg*), primers and probes for the single copy control gene (*nda2*, See Table S2 for oligos), 0.01 ng of genomic DNA, and the restriction enzyme MseI. We incubated the reaction mixtures at room temperature for 15 minutes to allow MseI digestion of the genomic DNA prior to droplet generation. We quantified the probe signal at the end of the ddPCR using Quantasoft software (BioRad). We calculated the standard deviation for each individual reaction using the formula

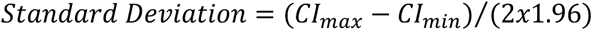

Where CI_max_- CI_min_ is the 95% Confidence Interval for the ratio of absolute copy number for *dg* and *nda2* in each reaction, with both assays multiplexed in the same well, as generated by Quantasoft. We calculated the copy number of the target sequence from the ratio of target sequence positive droplets to *nda2* positive droplets.

### Long-read sequencing and assembly of *S. pombe* wild isolates

We used an established protocol to extract high molecular weight genomic DNA from seven *S. pombe* strains (SZY44, SPT999a, SZY13, JB844, JB907, JB916, JB1172) (56). We prepared Nanopore sequencing libraries using Nanopore Ligation sequencing gDNA (SQK-LSK109). We sequenced the libraries on Nanopore Flongle (EXP-FSE001 and EXP-FLP002) or MinION (EXP-FLP002) flowcells.

We used Guppy (v6.4.6) to translate the raw Nanopore electrical signals into bases and explored the read quality with nanoplot (v1.33.0). To assemble the reads, we trimmed the sequencing adapters with porechop (v0.2.4), filtered the reads for quality and length using filtlong (v0.2.1; min_length=1000; keep_percent=90; target_bases=1380000000), then assembled the remaining reads using canu (v2.2; genomeSize=13.6m; correctedErrorRate=0.08). We polished the assembly by aligning the filtered reads to the assembly using bwa mem (v0.7.17), followed by one round of racon (v1.4.3; -m 8 -x -6 -g -8 -w 500) and medaka (v1.7.3) polishing. For JB907, centromere 3 did not fully assemble, so the Large centromere swap strain assembly (SZY5827; see below) was used for that centromere. We used centromeric sequence elements from the ASM294v2 reference genome (39) and BLAST searches to manually curate the sequence elements in Geneious Prime based on nucleotide alignments.

### Phylogeny Construction

We used MUSCLE (v3.8.1551) or Clustal Omega to align the annotated centromeric sequence elements and used the alignments to create phylogenies using PhyML (v3.3) with 1000 bootstraps. Any node with a bootstrap value >950 was deemed significant.

The entire centromere 3 of JB907 was not assembled (Fig S6A). Therefore, for the *dh* and *dg* sequence phylogenies, sequences from that centromere were replaced with the sequences from the Large swap strain (SZY5827; Table S1), whose centromere is derived from JB907 and was successfully assembled (Fig S6B). The rest of the sequences from JB907 are from the assembly of JB907 itself.

### Isogenic Strain Construction

Swapping distinct centromere 3 alleles into a common laboratory strain background involved a series of genome modifications and crosses as outlined in Fig S4. Key features included moving chromosomes intact between genetic backgrounds using *rec12Δ* crosses and using Cre recombinase to induce targeted recombination events flanking centromere 3.

### Construction of the adh1pr:LoxP:KanMX6 and LoxP:ura4:KanMX6 cassettes

We created these alleles as in (35) to create two plasmids, pSZB130 and pSZB132. These plasmids contained the needed alleles flanked by Hpa1 sites. We digested these plasmids and gel extracted and purified the cassettes. Those cassettes were then cloned into the Hpa1 site of pSZB1595, a construct synthesized by IDT. The Hpa1 site in pSZB1595 is in the middle of a 2kb region of homology to the right side of centromere 3 (centered around coordinate 1,142,476 on chromosome 3; reference genome ASM294v3) that is shared by the lab isolate, JB844, and JB907. This resulted in pSZB1614, which contains *adh1pr:LoxP:KanMX6* within the right cen3 sequence, and pSZB1662, which contains *LoxP:ura4:KanMX6* within the same sequence.

### Construction of the adh1pr:Lox2272:NatMX6 and Lox2272:his5:NatMX6 cassettes

We created these alleles as in (35) to create two plasmids, pSZB1131 and pSZB1308. These plasmids contained the needed alleles flanked by BamHI sites. We digested these plasmids, gel extracted and purified the cassettes. Those cassettes were then cloned into the BamHI site of pSZB1378, a construct synthesized by IDT. The BamHI site in pSZB1378 is in the middle of a 2kb region of homology to the left side of centromere 3 (centered around coordinate 1,043,403 on chromosome 3; reference genome ASM294v3) that is shared by the lab isolate, JB844, and JB907. This resulted in pSZB1451, which contains *adh1pr:Lox2272:NatMX6* within the left cen3 sequence, and pSZB1391, which contains *Lox2272:his5:NatMX6* within the same sequence.

### Creation of JB844 Hybrid Haploid 1 Strain (Fig S4)

We crossed the lab isolate strain SZY3834 to the JB844 background strain SZY3909 and screened for progeny that were *lys1-, leu1-,* and *ade6-*/Hyg^R^, which represented chromosome 1 and 2 from the lab isolate and chromosome 3 from JB844. This resulted in SZY4009, our first hybrid haploid strain.

### Creation of JB907 Hybrid Haploid 1 Strain (Steps 1-3, Fig S4)

JB907 appears to be sterile under lab conditions, so we generated our first hybrid haploid strain via protoplast fusion. We generated protoplasts from the lab isolate strain SZY3828 and the JB907 background strain SZY3902 via Lallzyme MMX digest as published (57). We then proceeded with protoplast fusion as published (58), selecting for a diploid based on complementary markers. We then put that diploid through meiosis and screened for progeny that were *ura4-, leu1-* and *his5-,* and *ade6-*/Hyg^R^, which represented chromosome 1 and 2 from the lab isolate and chromosome 3 from JB907. However, this exact genotype was not recovered as JB907 contains a chromosome 2-3 translocation that would make a strain with only chromosome 3 from JB907 inviable (Fig S4A). Therefore, we recovered a strain with chromosome 2 and 3 from JB907 and chromosome 1 from the lab isolate (SZY5181). We then swapped out chromosome 2 in the next construction step.

### Right Arm Swap to generate Hybrid Haploid 2 (Steps 4-6, Fig S4)

We digested pSZB1614 and pSZB1662 with KpnI to release the centromere 3 region targeting cassette and transformed them into the strains to cross. Digested pSZB1662 was transformed into a strain of the lab background to create SZY3938. Digested pSZB1614 was transformed into Hybrid Haploid 1 strains to create SZY4054 and SZY5385, respectively. We then transformed pSZB155 (from Dr. Steven Henikoff’s lab), which is a plasmid containing *LEU2* (from *Saccharomyces cerevisiae*, which complements *S. pombe leu1*) and a Tet-inducible Cre recombinase into SZY3938, SZY4054 and SZY5385. We grew the strains to be mated (Fig S4, steps 4-5) to saturation overnight in YEL at 32° C. We then combined 500 µL of the strains to be mated, spun the cells down, washed in water, and resuspended in 100 µL sterile water. We then spread this mixture onto sporulation media with supplements (SPA-S) with 45 mg/L tetracycline and incubated for 3 days at 25°C to allow for mating and meiosis. We then selected for *ura4+* progeny, indicating recombination has occurred at the centromere 3 adjacent site. This produced half-swapped Hybrid Haploid 2 strains (Fig S4; SZY4147 for JB844, SZY5467 for JB907).

### Left Arm Swap to generate isogenic swap strains (Steps 7-9, Fig S4)

We then digested pSZB1391 and pSZB1451 with NotI to release the centromere 3 region targeting cassette and transformed them into the strains to cross. Digested pSZB1451 was transformed into the lab isolate background to create SZY4927. Digested pSZB1391 was transformed into the half-swapped Hybrid Haploid 2 JB844 and JB907 hybrids to create SZY4925 and SZY5681, respectively. We followed the same steps of transforming the Cre recombinase plasmid and crossing on SPA-S+ tetracycline and selected for *his5+* progeny, indicating recombination has occurred at the centromere 3 adjacent site. This produced our first complete swap strains (SZY4960-4970 for JB44, SZY5746-SZY5748 for JB907).

### Generating the Medium centromere swap strain

Although the “Medium” strain has the same centromere allele as the lab isolate, we still performed the same strain construction steps to ensure the markers near the centromere would be identical in all tested conditions. The intermediate strains used in this strain construction are included in Table S1.

### Verification of Strains

The final copy number of *dg* was confirmed in each of the strains via ddPCR (see above). The sequence identity was confirmed via short and long read sequencing (see below).

### Long-read sequencing and assembly of swap strains

We performed long-read sequencing on our swap strains (Small SZY5658, Medium SZY5521, and Large SZY5827) as above. We outsourced additional Oxford Nanopore and Illumina short read sequencing to Plasmidsaurus (Small SZY5658, Medium SZY5521, Large SZY5827, and Jumbo SZY7388).

### Preliminary swap strain assemblies (Table S6, Figs S2, S3, S5A-B)

We performed the assembly steps listed above to generate preliminary assemblies. We used these assemblies in the phylogenies in Figures S2 and S3 and the mummer alignments in Figure S5A-B since the wild isolates were also assembled using this method.

### Final swap strain assemblies (all other figures)

We translated the raw Nanopore electrical signals from our in-house MinION flowcells into bases with dorado (v0.9.1; trim=all), which incorporates an adapter trimming step. We assessed the quality of the resulting reads with nanoplot (v1.33.0), length and quality filtered with filtlong (v0.2.1; min_length=600; keep_percent=80),and then assembled the reads using canu (v2.2; genomeSize=12.6m; correctedErrorRate=0.08). We polished these assemblies by aligning the filtered reads to the assembly using bwa mem (v0.7.17; -K 10000000), followed by a round of racon (v1.4.3; -m 8 -x 6 -g 8 -w 500) and medaka (v1.7.3) polishing.

We performed a second round of polishing with Illumina short read data to the final swap strain assemblies (Small SZY5658, Medium SZY5521, Large SZY5827, and Jumbo SZY7388) to minimize false polymorphisms primarily caused by the high error rate in Nanopore data around homopolymer runs. We aligned the Illumina paired-end 150bp reads to the racon and medaka polished assembly using bwa mem (v0.7.17), polypolish filter and polypolish polish (v0.6.1). Finally, we polished the assemblies with pypolca (v0.4.0; --careful) as recommended in (59), for genomes where Illumina read depth is >25x (Small = 168x, Medium = 1879x, Large = 199x, Jumbo= 1079x). Pypolca aligns short reads to the assembly using bwa mem (v0.7.17), processes the alignment with samtools (v1.20) and calls variants with freebayes (v1.3.10).

### Estimating Large centromere strain *dh* and *dg* copy number

Both the preliminary and final assemblies of the Large strain (SZY5827) contained a gapless assembly of chr3, but with 36 and 41 *dh/dg* repeats respectively. To attempt to resolve this discrepancy, we first filtered the raw Illumina reads from each strain with fastp (v1.0.1) and then mapped to *dh* and *dg* reference sequences with bwa-mem2 (v2.2.1) in a custom bash script. The *dh* reference consisted of a 1,634bp region (largely identical to coordinates 1,130,717 to 1,132,351 on chromosome 3, reference genome ASM294v3) conserved in all *dh* repeats, and the *dg* reference consisted of a 3,081bp region (largely identical to coordinates 1,133,254 to 1,136,361 on chromosome 3, reference genome ASM294v3) conserved in all *dg* repeats. Both references only contain sequence that is unique to the *dh* or *dg*. We also mapped the reads to eleven single copy *S. pombe* genes as controls (*emc5, meu27, per1, ppc1, rpb1, yox1, oac1, mrp123, coq4, gls2, and rib5*). We excluded reads that were less than 30bp in length or with less than 60% identity to the reference. For each reference, we calculated the average per-base read depth, excluding the first and last 50bp of each reference sequence. We estimated the copy number for the *dh* and *dg* by dividing the *dh* and *dg* average per-base read depths over the mean of the eleven control genes’ average per-base read depth (Table S6).

### Detecting polymorphisms between the swap strains

We aligned the final polished assemblies (see above) to ASM294v3 (39) using minimap2 (v2.26). Next, we used the minimap2 command paftools.js in a custom bash script to generate a variant vcf file, which we sorted with bcftools sort (v1.16). We generated a gff3 file containing all the variant calls with a custom awk command from the sorted vcf file. We compared the four swap strain gff3 files to each other using a custom python script that detects variants unique to each strain (Table S8-S11). We extracted information and genomic context for each of these unique variants (Fig S5C) with an additional python script, and all polymorphisms within protein coding sequences (based on the ASM294v3 annotations) were examined manually in Geneious Prime (v2026.0.1) (Table S7).

### Recombination Assay

Since all our swap strains had *NatMX6* and *KanMX6* integrated near the centromere, we transformed in *HphMX6* to replace either drug marker. To do this, we digested pAG32 (60) with the NotI restriction enzyme and transformed the digested plasmid into our swap strains. The similarity of the MX repeats in these markers allows for efficient replacement of one marker with another.

We grew the strains to be mated (see Table S13 for strains) to saturation overnight in YEL at 32°C. We then combined 500 µL of the strains to be mated, spun the cells down, washed in water, and resuspended in 100 µL sterile water. We then spread this mixture onto sporulation media with supplements (SPA-S) and incubated for 3 days at 25°C to allow for mating and meiosis. We then scraped the cells off the plate and treated with glusulase and ethanol (61). We then germinated 100 µL of spores in 5 mL of YEL at 32° C with shaking for 24 hours. We then diluted the cells and plated on YEA-S to assay cell concentration and on YEA-S containing 200 µg/mL G418 (Kan) and 100 µg/mL ClonNat (Nat) to select recombinants. We counted colonies after 2 days of growth at 32° C. Since some of the colonies on YEA-S +G418 +Nat could be chromosome 3 disomes, we also replica plated those media onto YEA-S +200 mg/L Hygromycin B, since disomes would still have a copy of *hphMX6*, and recombinants would not. We subtracted the total number of colonies on YEA-S +Hyg from the number of colonies on YEA-S +G418 +Nat to determine the total number of recombinants (Table S13). We used ddPCR (described above) to measure *dh*/*dg* copy number of some recombinant progeny.

### Spot dilution assays

To test growth on a variety of media, we grew each strain to saturation in YEL overnight before diluting 1:10 and growing for an additional 3 hours to ensure cells were actively growing in log phase. We then measured the OD_600_ using a spectrophotometer and diluted each culture to a standard starting OD_600_, usually OD_600_ = 0.5. We then performed 4-fold serial dilutions in sterile water and spotted each dilution onto the appropriate media using a frogger and incubated plates at the appropriate temperature. We then monitored the plates for growth and imaged after approximately 3 days at 32°C, or longer at lower growth temperatures.

### RNA sequencing

We grew the strains (Small SZY7163, Medium SZY7165, Large SZY7167, Jumbo SZY7595) in triplicate to OD_600_ ∼0.5 in fresh YEL broth at 32°C prior to RNA extraction following a published protocol (62). We used the RNA Clean & Concentrator kit (Zymo Research) to clean up the RNA. We depleted the ribosomal RNA with FastSelect Yeast reagent (Qiagen) and prepared the RNA libraries with RNA-Seq FastSelect Depletion (stranded) (Watchmaker). We sequenced the libraries with the AVITI sequencer (Element Biosciences).

We demultiplexed the raw RNA reads into Fastq format allowing up to one mismatch using Illumina bcl-convert 3.10.5. We aligned the reads to UCSC genome ASM294v2 (39) with STAR aligner (version 2.7.11b), using Ens_fungi_57 gene models. We generated TPM values using RSEM (version v1.3.1) and we used Bioconductor package edgeR (3.24.3 with R 3.5.2) to perform pairwise differential expression analysis. We considered only protein coding genes and long non-coding RNAs (lncRNAs) from the Ens_fungi_57 annotation. We considered only genes with counts per million expression of at least 0.5 in at least two samples for further analyses. We determined statistical significance by fold change cutoff of 2 and false discovery rate (FDR) cutoff of 0.05.

### Measuring chromosome 3 disomy

Strains to be mated (see Table S16 for strains) were mated as in the recombination assay above. We plated the glusulase and ethanol treated spores on rich medium (YEA-S) for 6 days at 32°C to allow germination.

We then picked colonies from YEA-S plates with fewer than 300 colonies and patched them onto a master plate (YEA-S) that was grown for an additional 2 days at 32°C. We then replicated the master plates onto 1) YN-ade plates 2) YEA-S + 200 mg/L Hygromycin B and 3) YEA-S plates. We grew the replica plates for two days at 32°C before scoring. We called a colony *ade6+* if any part of the patch grew white on YN-ade, and Hyg^R^ if any part of the patch grew on YEA-S +Hyg. The growth was often inconsistent within a patch as chromosome 3 aneuploids are unstable.

### Timelapse Microscopy

We introduced fluorescent reporters into the swap strains (Small SZY7163/7164, Medium SZY7165/7166, Large SZY7167/7168, Jumbo SZY7595/7597) by crossing in the Hht1-RFP marker from an existing strain (SZY968) (63). Then, we digested plasmid pAde6PmeI-patb2-sfGFP-Atb2-terminatoratb2-hphMX (64) with PmeI and integrated it into the a*de6* locus.

We grew each strain (Small SZY7170, Medium SZY7174, Large SZY7179, Jumbo SZY7611) to saturation at 32°C in YEL media overnight. We then spotted 50 µL of each strain onto both YEA-S + 1% DMSO and YEA-S + 20ug/mL TBZ media and grew for 24 hours at 32°C. We then scraped the cells off the plates into 100 µL of sterile water and then spread the cells mixture onto a fresh plate of the same type with a sterile pipette tip. We then cut agar punches from these plates and inverted them into a coverslip bottom multiwell chamber slide. We imaged the cells on a Nikon Ti microscope coupled to a spinning disc (Yokogawa CSU-W1). We kept the cells were 32°C with a stage top incubator (Oko Lab). We excited the GFP and RFP tagged proteins at 488 and 561 nm, respectively through a 100x Plan Apo objective (NA 1.45). We collected transmitted light images and fluorescent images through ET535/30m (GFP) and ET605/70m (RFP) filters every 20 minutes with z-steps of 0.5 μm through a total of 5 μm. The still images shown in Fig 4B are max-projected across all z-steps.

To measure symmetry of divisions, we drew regions of interest by hand in Fiji (https://imagej.net/software/fiji/downloads) around recently divided cells where the septum had begun to form, but cytokinesis had not yet been completed. We then loaded those regions of interest into a custom written python notebook along with the raw data. We sum projected the images and identified the dividing nuclei. We discarded all cells with more than two objects (lagging chromosomes, etc.). We then found the sum intensity of each nucleus, and the lower intensity of the two was divided by the higher intensity nucleus to determine the intensity ratio plotted in Fig 4C.

## Supplemental Information Legends

**Table S1. Strains.** All strains used in this study are listed with their full genotypes. Strains assayed in figure S1 come from a previously published natural isolate collection (33).

**Table S2. Oligos. The** oligos used for ddPCR of each sequence element are listed. ddPCR probes contain a 5’ fluorescent probe (HEX or FAM) a spacer (ZEN) and a 3’ quencher (3IABkFQ).

**Table S3. Plasmids.** All the plasmids involved in strain construction and marker creation are listed.

**Table S4. Natural Isolate ddPCR screen.** *dh* and *dg* copy numbers of 161 natural isolates were screened via ddPCR. Copy numbers of all 3 replicates are listed. NA indicates that there were insufficient droplets to calculate a copy number in that replicate. Strains assayed in this analysis originated in (33). Copy number was determined by dividing *dh* or *dg* positive droplets by the number of *nda2* positive droplets.

**Table S5. Repeat Sequence Identity.** Identity of each sequence element compared to the consensus sequence of all elements.

**Table S6. Large Copy Number (CN) Estimates.** Different assembly methods (dorado (v0.9.1; trim=all), or guppy (v6.4.6) basecalling) yielded either 36 or 41 copies of *dh* and *dg* on chromosome 3 of the Large centromere strain, but the Small and Medium strains remained consistent. Jumbo centromere 3 was never fully assembled (top). We used high coverage Illumina sequencing to calculate the mean read depth of *dh* and *dg* in each strain. We used the coverage of 11 different single-copy control genes to estimate the single gene copy number. The read depth for each control is shown alongside the *dh* or *dg* copy number estimate (repeat coverage/single gene coverage) for each (middle). The average read depths across all 11 single-copy genes are shown (bottom).

**Table S7. Coding Sequence Variants.** All variants found in the Small, Medium, Large, or Jumbo strain that fall within coding variants when aligned with the Pombase reference sequence (ASM294v3) are shown. For nonsynonymous mutations, the amino acid changes are shown.

**Table S8. Small Centromere Strain Variants.** All variants found in the Small centromere strain when aligned with the Pombase reference (ASM294v3) are shown.

**Table S9. Medium Centromere Strain Variants.** All variants found in the Medium centromere strain when aligned with the Pombase reference (ASM294v3) are shown.

**Table S10. Large Centromere Strain Variants.** All variants found in the Large centromere strain when aligned with the Pombase reference (ASM294v3) are shown.

**Table S11. Jumbo Centromere Strain Variants.** All variants found in the Jumbo centromere strain when aligned with the Pombase reference (ASM294v3) are shown.

**Table S12. Recombination Frequency.** Data plotted (top) was calculated by counting total colony forming units (CFU) based on colony counts on YEA-S and recombinant CFU based on colony counts on YEA-S +Nat +G418 minus the colonies which grew on YEA-S +HYG (disomes). The raw colony counts (bottom) are shown for each plate in each cross.

**Table S13. Large Recombinant Screening.** DNA was isolated from Large strain progeny which appeared to be changing in copy number based on an initial screen (left). Note that the copy number assayed in most progeny appeared to be very close to that of the Large parents. Known Large (SZY6581, SZY6582, and SZY6615) and medium strains (SZY7044, SZY7045) were also included concluded as controls. We discarded any reactions with an *nda2* copy number of less than 8. One strain, SZY7155, appeared to be larger than its parent, so this strain was retested, starting with a fresh DNA preparation (right). For all samples, 3 replicates are shown with *dg* and *nda2* counts for each.

**Table S14. Mitotic Stability ddPCR.** We isolated DNA from Small, Medium, Large, and Jumbo strains before and after eight days of continuous mitotic growth. The plotted data (top) is the average of three technical replicates (bottom) of one independent colony. We discarded any reactions with a *nda2* copy number of less than 8 (highlighted in red).

**Table S15. All RNA-Seq hits**. Tables list all differentially expressed genes (fold change counts per million >2, false discovery rate <0.05) in the Small (top), Large (middle), and Jumbo (bottom) centromere strains when compared to the Medium centromere strain.

**Table S16. Chromosome 3 Disomy.** We picked colonies from crosses of the indicated centromere sizes and replica plated to determine disomy state. Percentage of Disomy (top) was calculated with at least three independent replicates (bottom) for each cross.

**Table S17. Division Symmetry Measurements.** For each timelapse, we drew regions of interest (ROIs) around each set of cells with recently divided nuclei. We calculated the sum signal intensity each nucleus within the ROI, calculated the ratio of signals by dividing the smaller intensity by the larger intensity.

## Acknowledgements

We thank the members of the Gerton and Zanders lab for discussions of the work. We thank Dr. Harmit Malik and Dr. Mia Levine for their insights on our work. We thank Dan Bradford for his help in developing the ddPCR screening methodology. We thank Dr. Sergio Tusso for sharing the dendrograms of the natural isolates. We thank Dr. Shiv Grewal for sharing the SPT999a strain. We thank Dr. Steven Henikoff and Dr. Takehito Furuyama for sharing the inducible Cre plasmid. Deletion strains to create mutants were provided by the NBRP, Japan (JPNBRP202225).

## Data Availability

Whole genome sequencing and RNA sequencing data is available at the NCBI SRA under accession number PRJNA1428304. Original data underlying this manuscript can be accessed from the Stowers Original Data Repository at https://www.stowers.org/research/publications/LIBPB-2612. Relevant code can be found on the Zanders-Lab GitHub page.

---

**Figure S1.**
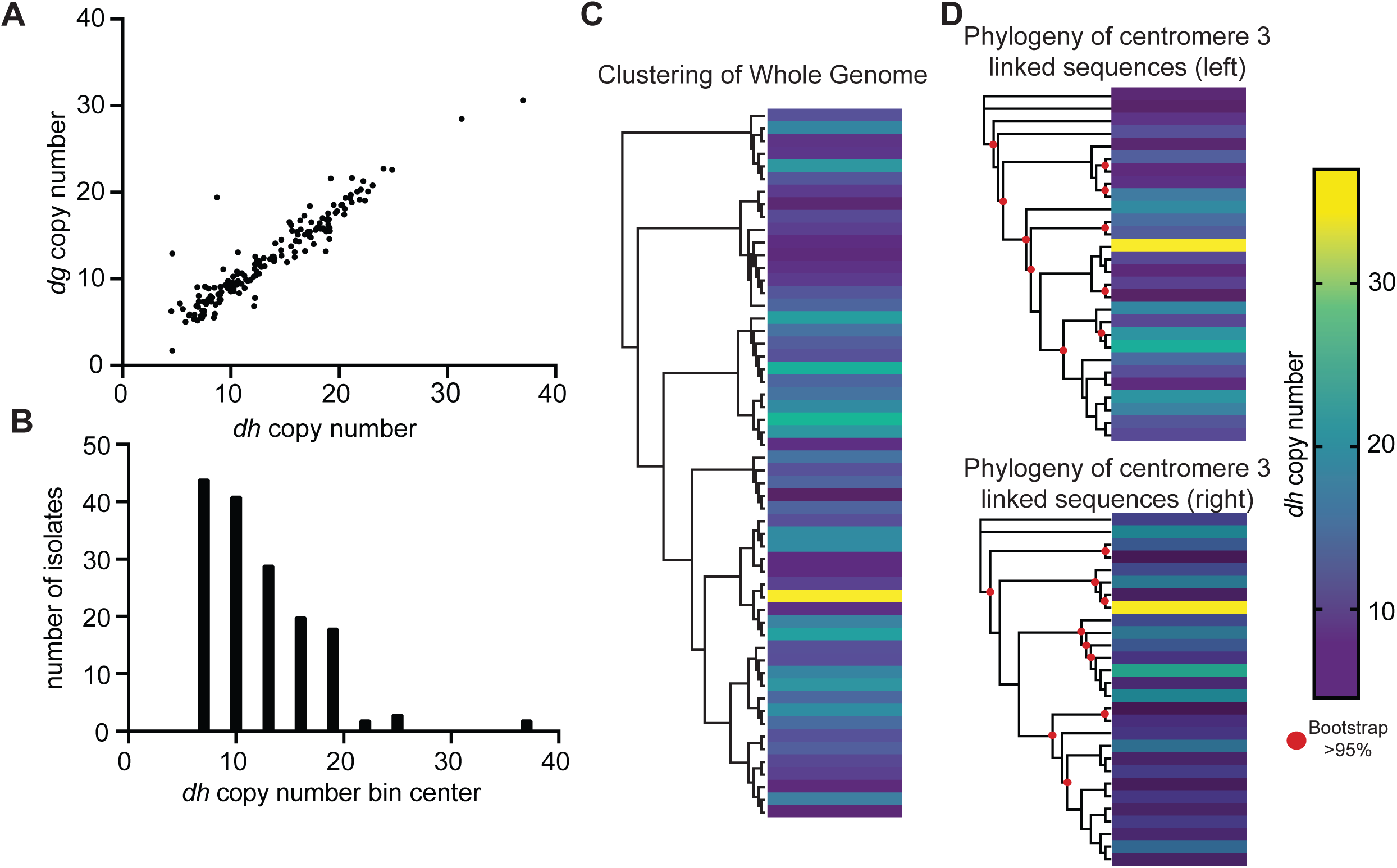
*dh* and *dg* copy numbers are rapidly co-evolving. A) The copy number of *dh* and *dg* repeats found in 159 natural isolates assayed by droplet digital PCR (ddPCR) is plotted, showing a strong correlation between the two repeats. Means of 2-3 technical replicates are shown. All technical replicates are shown in Table S4. B) Histogram of frequency of each *dh/dg* copy number from the set of 159 isolates measured. Means of 2-3 technical replicates are plotted in bins centered around 7, 10, 13, 16, 19, 22, 25, and 37. C) Dendrogram constructed using whole genome sequence data of 56 non-identical natural isolates from (65). For sequenced isolates that are not represented in the dendrogram, their clonal derivative (33) or closest relative is shown. Color coding is based on the *dh* copy number of each isolate measured by ddPCR, as shown in the legend. D) Phylogenies constructed using 30kb of sequence directly upstream (top) or downstream (bottom) of centromere 3 from 28 sequenced isolates (Clustal Omega alignment, PhyML 1000 bootstraps). Color coding is based on the *dh* copy number of each isolate measured by ddPCR, as shown in the legend.

**Figure S2.**
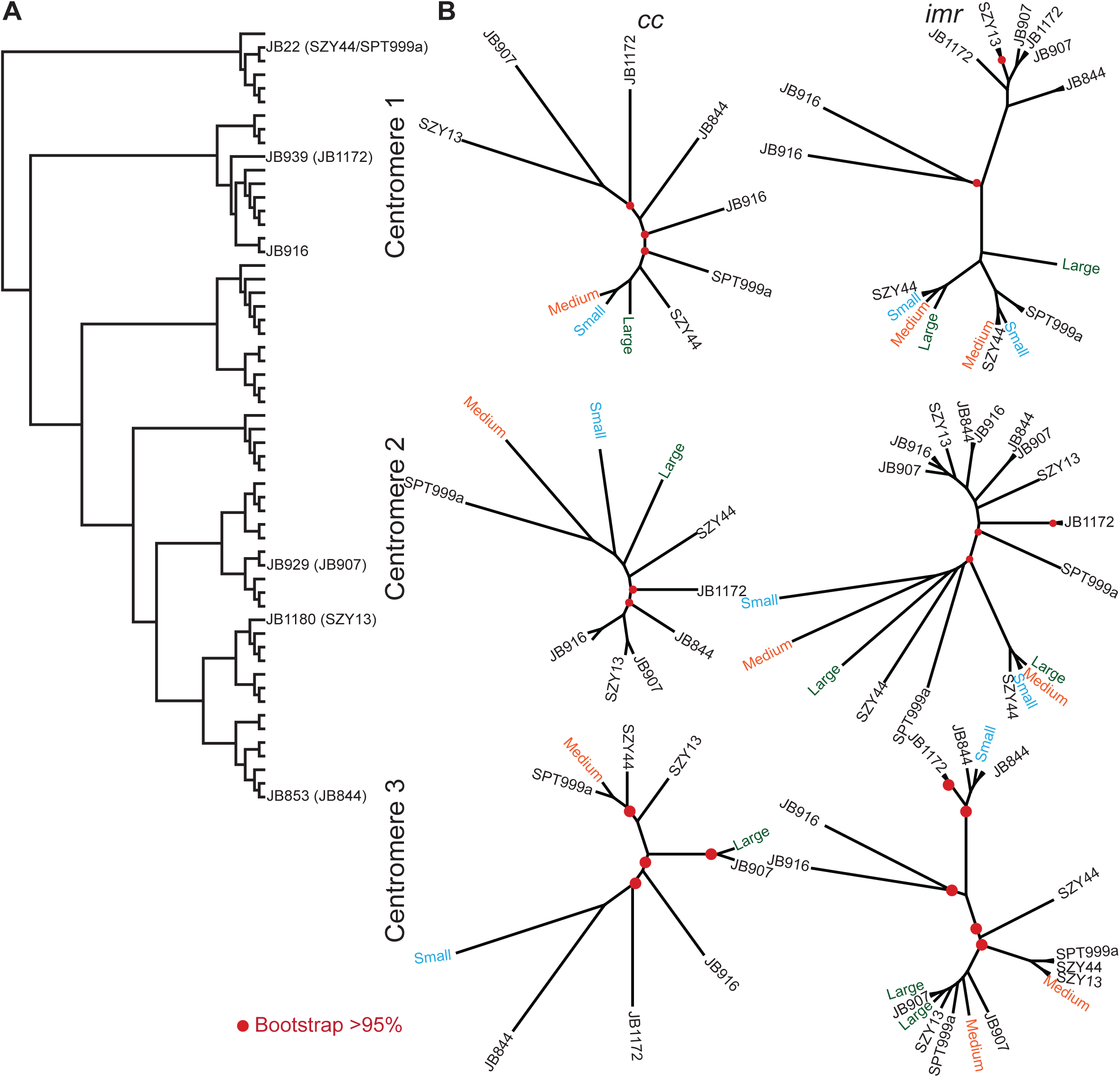
Phylogeny of defined centromere regions from the assembled centromeres. A) The relationship between the natural isolates used in this study and other *S. pombe* natural isolates is depicted in the dendrogram (65). Isolates are arranged from *S. pombe* like (top) to *S. kambucha* like (bottom) (65). For sequenced isolates that do not appear in the dendrogram their clonal derivative or closest relative is shown, with the sequenced isolate used in our study in parentheses (33). B) Maximum likelihood trees (PhyML with 1000 bootstraps) of the central core (*cc*; left) and innermost repeat (*imr*; right) sequences of centromeres 1-3 (top to bottom) from the seven assembled isolates and 3 swap strains (Fig 3A). Branches with >95% bootstrap support are indicated with red dots. Note that JB916 has a duplication of centromere 2, resulting in 2 *cc2* sequences and 4 *imr* sequences.

**Figure S3.**
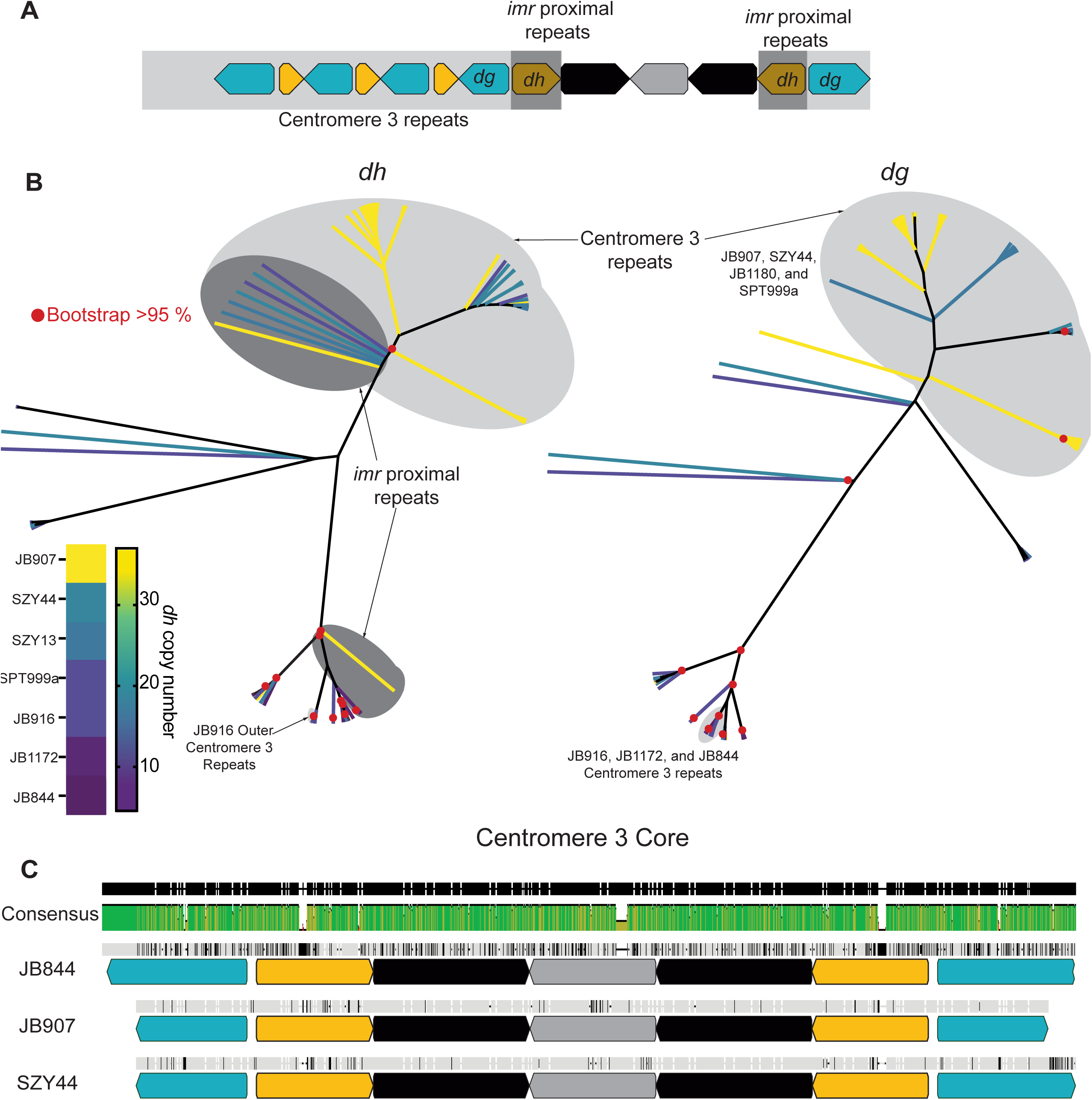
*dh* and *dg* sequence variation. A) Schematic of centromere 3 illustrating the position of the repeats used in the phylogenies shown in B. B) Maximum liklihood tree of all dh (left) and dg (right) repeats from all chromosomes assembled in this study (PhyML, 1000 bootstraps). Each branch represents one repeat unit and nodes with >95% bootstrap support are indicated with red dots. The branches for the other repeats found on centromere 3 are shaded in light gray. The branches for dh repeats adjacent to the *imr3* repeats are shaded in dark gray. The branches are color coded as shown in the legend (bottom left) based on strain/total *dh* copy number. C) Alignment of the core of centromere 3 in three natural isolates shows 95.4% sequence identity.

**Figure S4.**
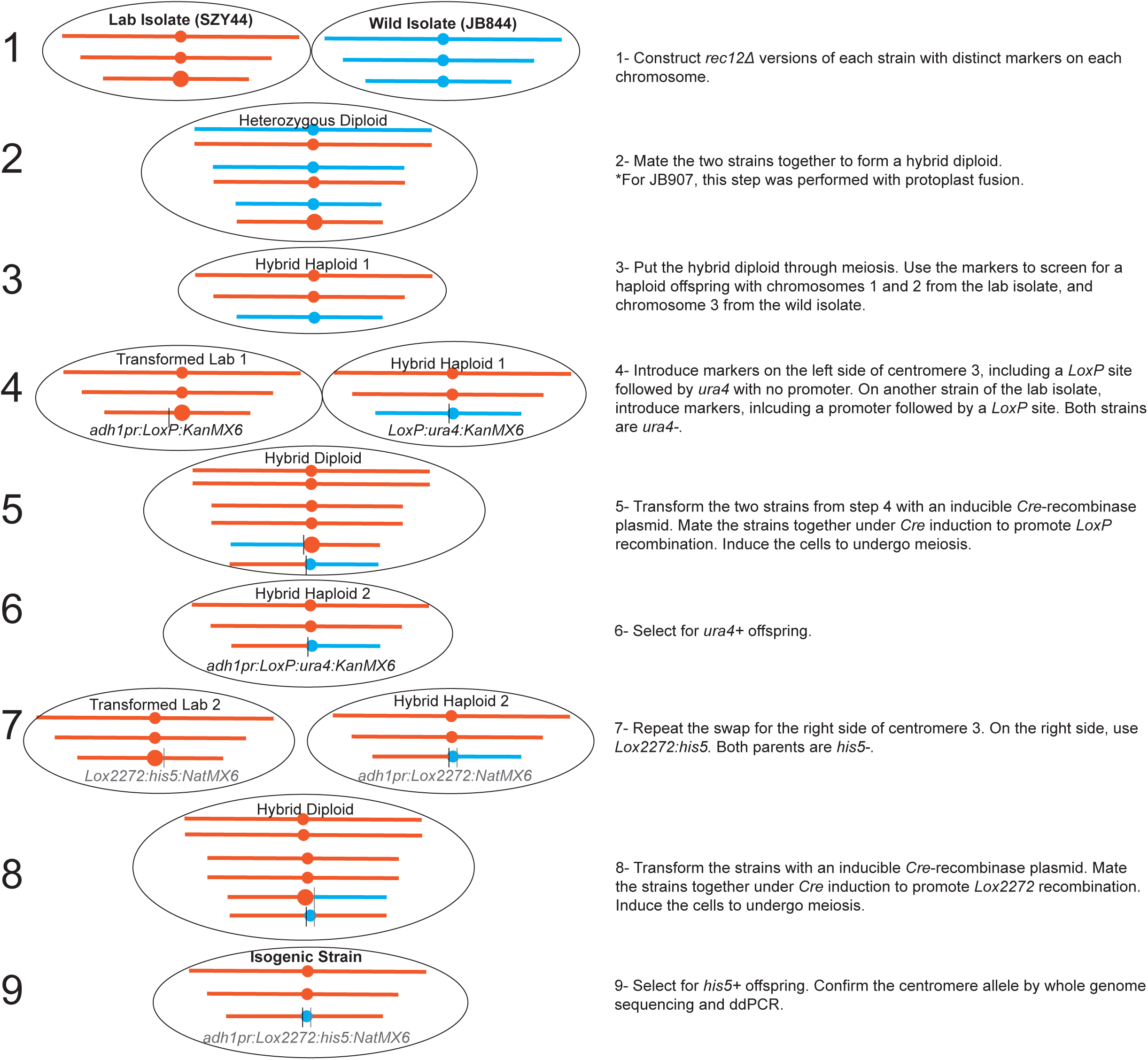
Construction of isogenic centromere 3 variant strains. The steps taken to construct the isogenic centromere 3 variant strains are described in 9 steps. The strain names are available in Table S1 and the plasmids and constructs are in Table S3.

**Figure S5.**
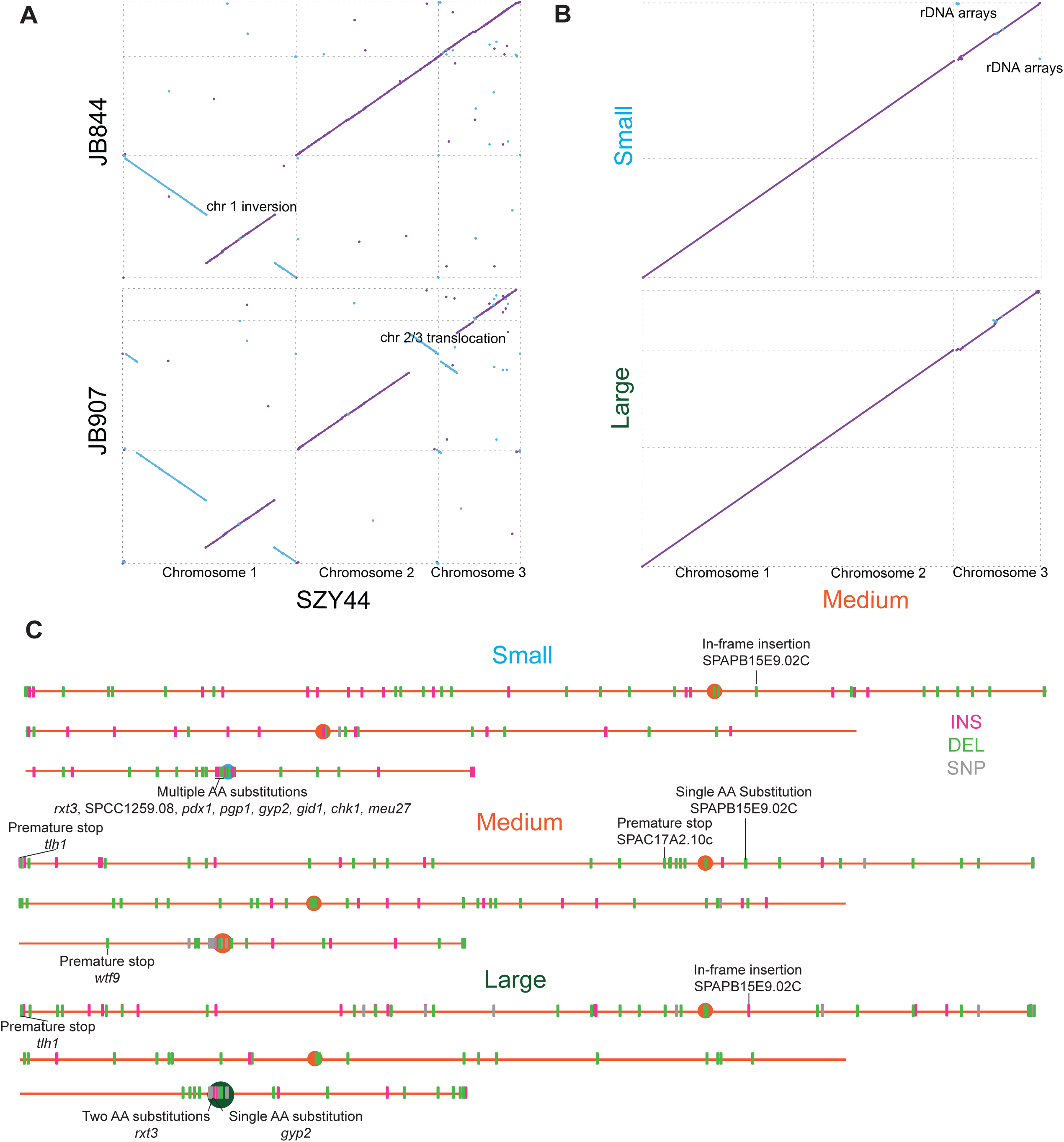
Centromere 3 variant strains are isogenic. A-B) Alignments of the natural isolates that were used to generate the isogenic centromere 3 swap strains (A) and the swap strains themselves (B) are shown as dotplots. The alignments and plots were generated by MUMmer. Cyan diagonals represent inversions, purple diagonals represent sequences in alignment. C) Location of all unique variants (insertions (INS), deletions (DEL), or single nucleotide polymorphisms (SNP)) located in the 3 swap strains. All non-synonymous variants within coding regions are labeled in black. All variants listed in Tables S7-10.

**Fig S6.**
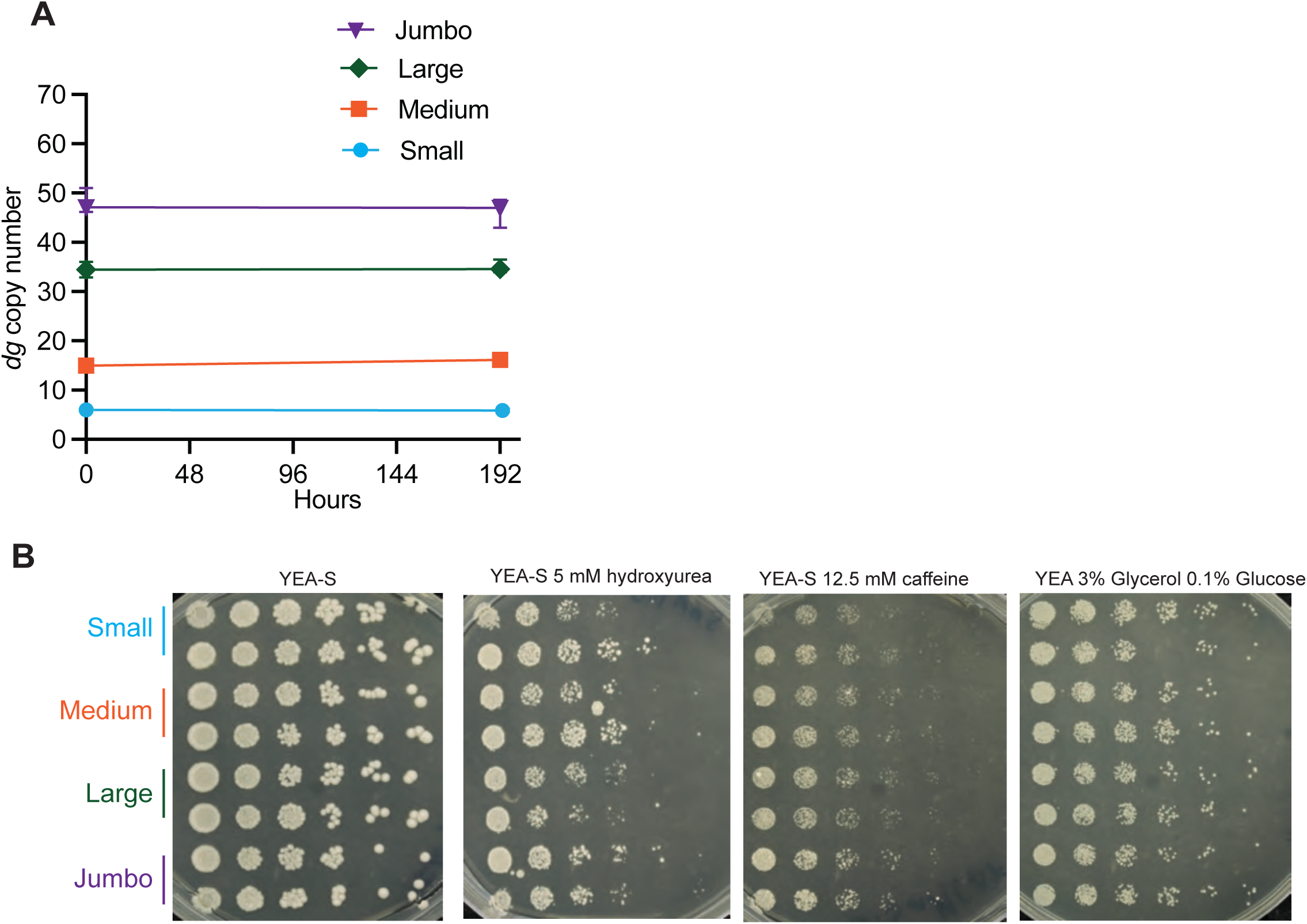
Isogenic strains stably maintain their copy number and show similar stress tolerance. A) *dg* copy number of the isogenic strains, measured by ddPCR, before and after 8 days of continuous mitotic growth in rich liquid media at 32°C, with saturated cultures diluted 1:1000 in fresh medium every 24 hours, exceeding the amount of doublings required for a standard spot assay. Raw data shown in Table S14. B) Spot assays of Small, Medium, Large, and Jumbo cen3 strains (2 biological replicates shown) were performed on indicated media. All plates were incubated for 3 days at 32°C prior to imaging.

**Figure S7.**
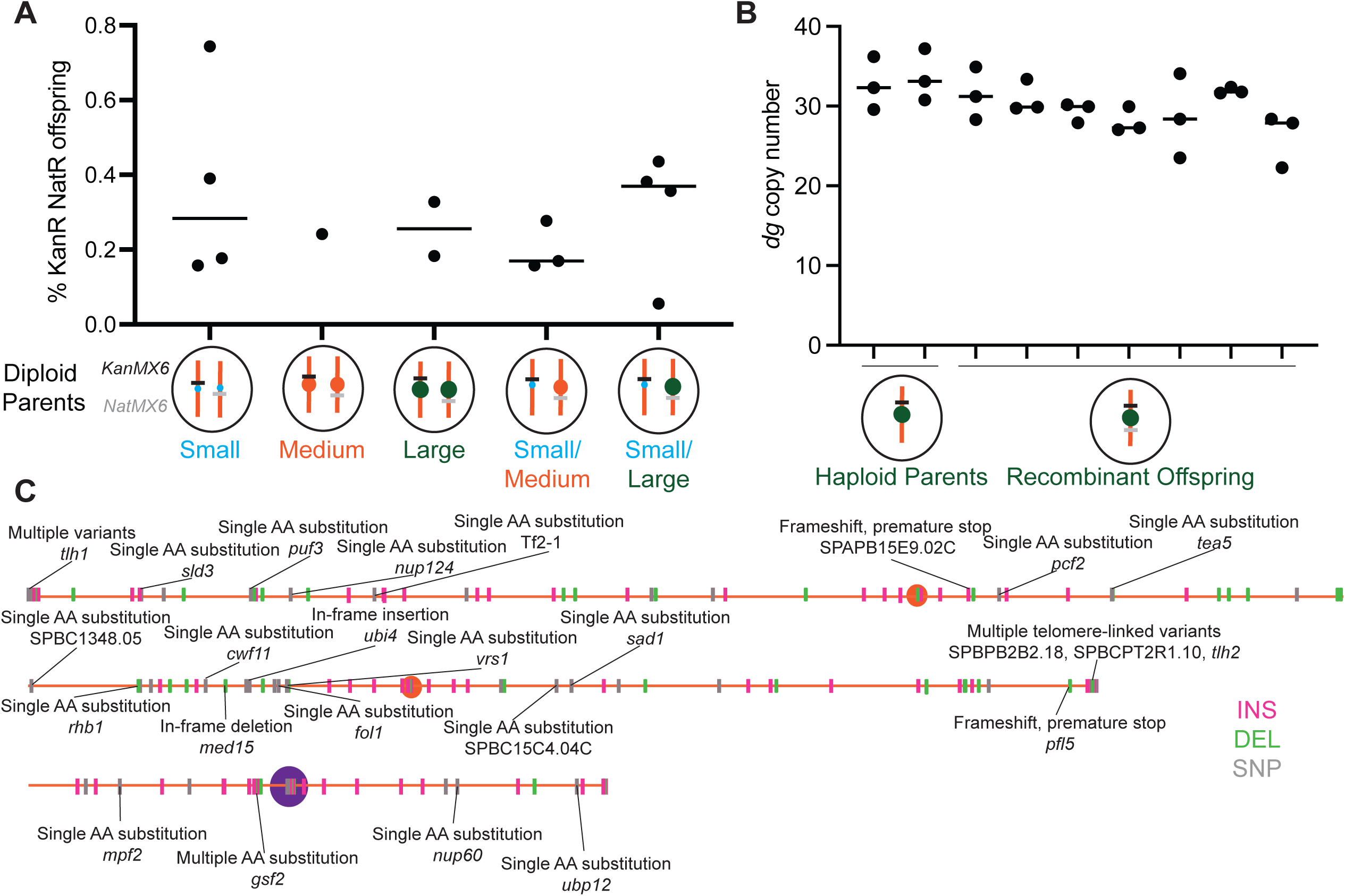
Crossover events between centromere-linked markers are rare and unaffected by *dh/dg* copy number. A) The percentage of progeny which have undergone a crossover between centromere adjacent markers in diploid parents with the given genotype, along with the median values (lines) is shown. Each point represents one biological replicate of the cross with at least 100 recombinants counted. The full counts are available in Table S12. B) *dg* copy number of the haploid parental Large strain and a subset of progeny which have undergone a crossover between centromere adjacent markers. The copy number was measured by ddPCR in technical triplicate. All three technical replicates are plotted, along with the median values (lines). A subset of progeny is shown. The raw data are available in Table S13. C) The location of all unique variants (insertions (INS), deletions (DEL), or single nucleotide polymorphisms (SNP)) located in the newly created “Jumbo” strain is shown. All non-synonymous variants within coding regions are labeled in black. A full list of variants is available in Table S11.

**Figure S8.**
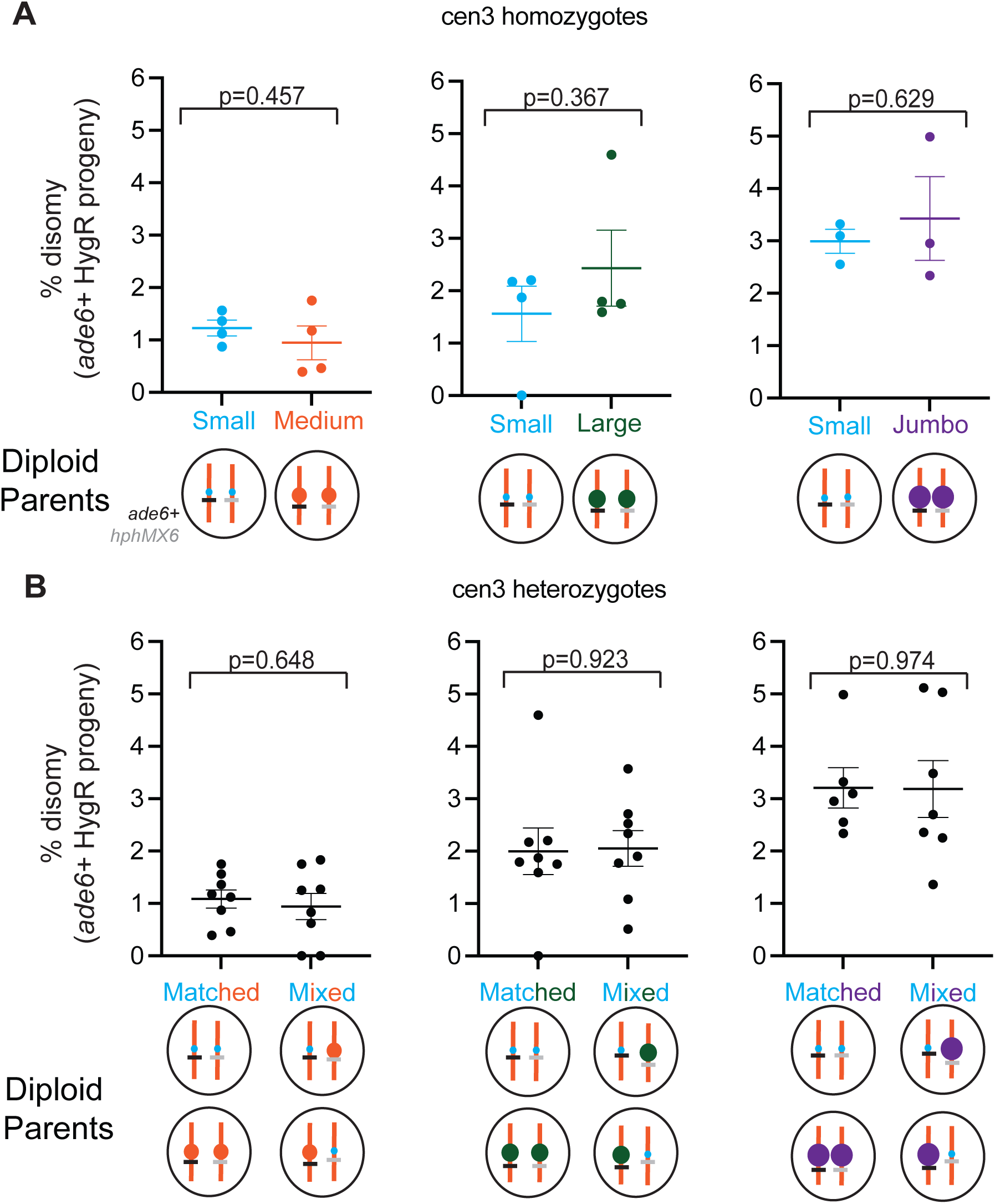
Chromosome 3 disomy is unaffected by pericentromere 3 array size. A) The fraction of disomic spores produced by diploids homozygous for Small, Medium, Large, or Jumbo centromeres is shown. Each dot is a biological replicate with n>300 spores genotyped. The lines depict the mean +/- SEM across biological replicates. p-values were calculated by an unpaired t-test. B) The fraction of disomic spores produced by diploid parents homozygous and heterozygous for centromere size is shown. Each dot is a biological replicate with n>200 spores genotyped. p-values were calculated by an unpaired t-test. Full counts are available in Table S16.

**Figure S9.**
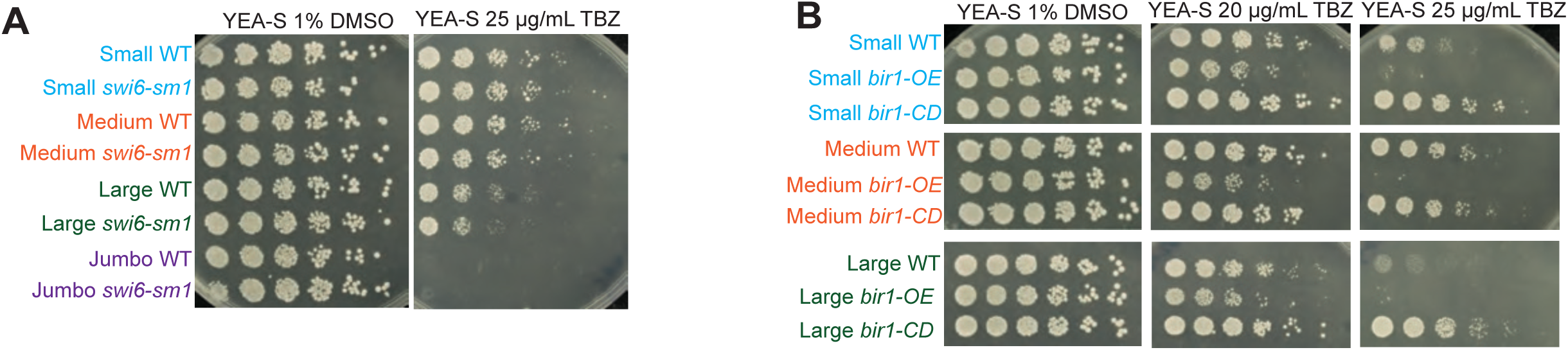
Heterochromatin and CPC factors modulate TBZ sensitivity. A-B) Spot assay of the indicated genotypes and size alleles are shown. Note that in panel B, white lines between plates indicate the strains were not grown on the same plate.

